# Heterologous immunization modulates B-cell epitope competition between helper peptides and the MPER segment in MPER/liposome vaccines

**DOI:** 10.1101/2025.04.23.650289

**Authors:** Rafiq Ahmad Khan, Junjian Chen, Luke Donius, Ellis L. Reinherz, Mikyung Kim

## Abstract

Subdominant B-cell immune responses to conserved epitopes are major obstacles in eliciting broadly neutralizing antibodies (bnAbs) against HIV-1 through natural infection or vaccination. Although the sequence conserved membrane proximal external domain (MPER) of HIV-1 gp41 is partially occluded on the virion surface, epitope-focused immunogens could mitigate access limitations. Here, we found that a MPER/liposome vaccine delivered with single CD4 T cell helper epitope results in a post-priming response hierarchy, eliciting low affinity MPER-specific B cells. Heterologous boosting, however, promotes MPER-specific B cell clonal expansion and enhances plasma antibody functionality. This improvement is associated with increased B cell affinity for MPER and reduced competition from B cells targeting the helper epitope. While helper peptide co-delivery increases affinity of serum antibodies, the outcome of subsequent MPER antibody responses is shaped by the priming antigen. Our results offer insights into heterologous immunization strategies to potentiate subdominant B cell responses against frequently mutating viruses.

**Significance Statement:** A key challenge in vaccination against mutable viruses like HIV-1 is the immune system’s focus on highly variable regions. The conserved MPER segment of gp41 elicits weak antibody responses due to poor B cell accessibility. This study evaluated a liposome-based MPER vaccine strategy. Initial priming generated low-affinity MPER-specific memory B cells, influenced by strong B cell affinity for a dominant T cell helper peptide. However, heterologous boosting overcomes subdominant MPER responses by increasing B cell affinity for booster immunogens while also reducing competition from other helper peptides. Co-delivery of helper peptides enhanced antibody affinity, though the priming antigen was critical in shaping responses. These findings suggest heterologous immunization is a promising strategy to enhance subdominant B cell responses.

## Introduction

Vaccines against viruses provide protection primarily by inducing neutralizing antibodies (nAb) that prevent the acquisition of infection ^1,2^. However, rapidly mutating viruses such as HIV-1 can evade host humoral responses through viral diversification ^3-5^. In such cases, it is of great interest to develop vaccination strategies to elicit neutralizing antibodies against conserved epitopes that are not under immune pressure. Antibodies recognizing these epitopes can be cross-reactive and confer broad protection, but no effective vaccine against the highly diverse viral strains is yet available ^6-8^.

A subset of people with HIV-1 develop potent and broadly neutralizing antibodies (bnAbs) within 2-3 years of infection, whereas the majority of people with HIV-1 generate non-neutralizing or strain-specific nAbs ^9-12^ preferentially targeting immunodominant variable domains in the envelope (Env) surface spike protein, the gp160 trimer. Structural and genetic characterizations of bnAbs developed from people with HIV-1 show unusual characteristics such as extensive somatic hypermutations (SHM) ^13,14^, highly improbable functional amino acid mutations ^15^, rare bnAb B cell precursors and autoreactivity ^16,17^. The unusually high rates of SHM indicate an extensive B cell affinity maturation process necessary for most antibodies to achieve effective HIV-1 neutralization. In addition, B cell receptor (BCR) access to the conserved epitopes on Env gp120 is shielded by glycans estimated at greater than 50% of the molecular mass of the Env ^18^, ^19^. Meanwhile, access to the neutralizing target on the gp41 subunit referred to as the membrane proximal region (MPER) is hindered by its location proximal to the viral membrane ^20-24^. These intrinsic biophysical properties of the Env may profoundly impact B cell competitive fitness, giving rise to immunologically subdominant responses against the conserved neutralizing epitopes.

Through biophysical, structural and genetic characterizations of recombinant bnAbs, reverse vaccinology has provided critical insights on the new vaccine concepts such as structure-guided and epitope-focused immunogen design as well as germline targeting immunogen design. These approaches aim at inducing dominant antibody responses toward the conserved epitopes such as a quaternary site at the V2 trimer apex, a V3-glycan gp120 supersite, the fusion peptide, the MPER or the CD4 binding site ^6,25^, shifting away from the majority of strain specific-antibody responses. While germline-targeting immunogens with increased antigen valency and/or affinity for bnAb precursors serve as priming antigen to expand bnAb precursor B cells, designing multiple boosting immunogens and implanting effective sequential immunization strategies remains a significant challenge in the pursuit of affinity maturation and bnAb development of the primed bnAbs precursors^26-31^.

With respect to the various epitope-focused vaccine targets, the MPER is amongst the most conserved across diverse HIV-1 clades and strains. That conservation and the broadest neutralization breadth manifest by MPER-specific bnAbs ^25,32-40^ warrant its contiguous structural epitopes as key targets for vaccine design. However, its membrane proximal location at the base of Env trimer partially embedded in virion membrane hinders naïve B cell access, resulting in immunologically poor responses elicited by vaccination with membrane-bound Env trimers ^41^. Many bnAbs exploit hydrophobic residues at the apex of their long heavy chain complementarity-determining region 3 (CDR3H) loop to facilitate interaction with the MPER and vicinal lipids to achieve neutralization breadth and potency ^42-49^. In that regard, a high density MPER segment array on the surface of liposomes provides a membrane environment necessary for the elicitation of bnAb paratopes. Accordingly, the epitope-focused MPER/liposome vaccines resulted in eliciting high titer anti-MPER antibodies ^50,51^. However, predominantly non-neutralizing sera Abs were elicited in a murine model, unable to bind the MPER of gp160 embedded on a membrane surface due to the incomplete quaternary structural mimicry of the MPER/liposome immunogen ^49^. Nonetheless, a recent clinical human trial study with MPER/liposome vaccine observed that although there is no serum neutralizing activity per se detected, monoclonal antibodies rescued from rare B cells in immunized volunteers did exhibit some cross-reactive neutralizing activity ^51^. This proof-of-concept emphasizes a critical need for developing vaccine strategies to further expand those B cells and gain sufficient neutralizing activity of serum antibodies.

Studies suggest that a heterologous vaccination regimen confers better protection against SARS-CoV-2 ^52-54^. Boosting strategies are important to provide additional antigenic stimulation, facilitating the refinement of the antibody repertoire to enhance neutralization potency and breadth ^55,56^. Likewise, a successful HIV-1 vaccine regimen will likely require sequential vaccination employing various antigens to expand and affinity mature the potential bnAb precursors specific to HIV-1 Env ^57^. Given that immunodominance is dependent on the antigen context ^58^, factors that can modulate antibody responses through boosting immunogens are not well understood, particularly in the heterologous vaccination setting. Using a limited number of B cell epitopes in the context of MPER/liposome vaccines as bnAb targeting immunogen models, here we explored booster immunization strategies. We evaluated how subdominant MPER-specific B cell responses are modulated after priming by competing B cell epitope responses directed at helper peptides following the subsequent boosting. While heterologous antigen boosting increases the anti-MPER antibody titers, the degree of plasma cell expansions and affinity maturation of the MPER-specific antibodies are greatly influenced by differences in affinity of B cells for the target MPER versus the competing helper peptides as well as the co-delivery of CD4 T cell helper peptides.

## Results

### pMPERTM/liposome vaccine elicits a dominant antibody response directed to LACK helper peptide antigen

Previously, N- or C-terminally palmitoylated (p) MPER segments (referred to as NpMPER or CpMPER, a.a 662-682 of HxB_2_ gp160) arrayed on the surface of a liposome vaccine induced strong MPER-specific antibody responses compared to gp160 immunogens. However, while anchoring the MPER to the membrane via palmitic acid or the transmembrane domain of the HIV-1 gp41 subunit facilitated co-delivery of the MPER segment and liposome vaccine components, the MPER chemical conjugation strategies influenced MPER membrane orientation and antigenic residue accessibility of the same MPER sequence, and consequently, resulted in distinct epitope specificities of the elicited antibodies by those various MPER immunogens derived from HxB_2_ strain^59^. For example, NpMPER/liposome elicited a dominant antibody response directed to the C-terminal region of the MPER involving a key epitope residue W680 and the chemical adduct NH_2_. On the other hand, using a liposome vaccine incorporating an immunogen where the MPER preceded the transmembrane domain (MPERTM) induced a majority of antibody responses specific to the N-terminal region of MPER including an acetyl group (CH_3_CO) at the N-terminal end (Supplementary Fig. 1A).

Accordingly, as shown in Fig. 1A we synthesized an N-terminally palmitoylated MPERTM (pMPERTM) immunogen to eliminate undesirable BCR interaction with the NH_2_ or the CH_3_CO chemical adduct at the C-and the N-terminal end and investigated its immunogenicity in a pMPERTM/liposome formulation. Balb/c mice were immunized with either pMPERTM/liposomes (G1) or with NpMPER/liposomes (G2), for comparison. Both liposome vaccine formulations incorporated the soluble LACK MHC class II binding (I-A^d^) helper peptide aa 156-173 (sLACK) derived from the Leishmania parasite^60^ along with a cyclic dinucleotide (CDN) adjuvant, specifically cyclic diadenylate monophosphate (c-di-AMP), which acts as a STING agonist ^61^. Both components were equivalently encapsulated inside the liposomes.

**Fig. 1.**
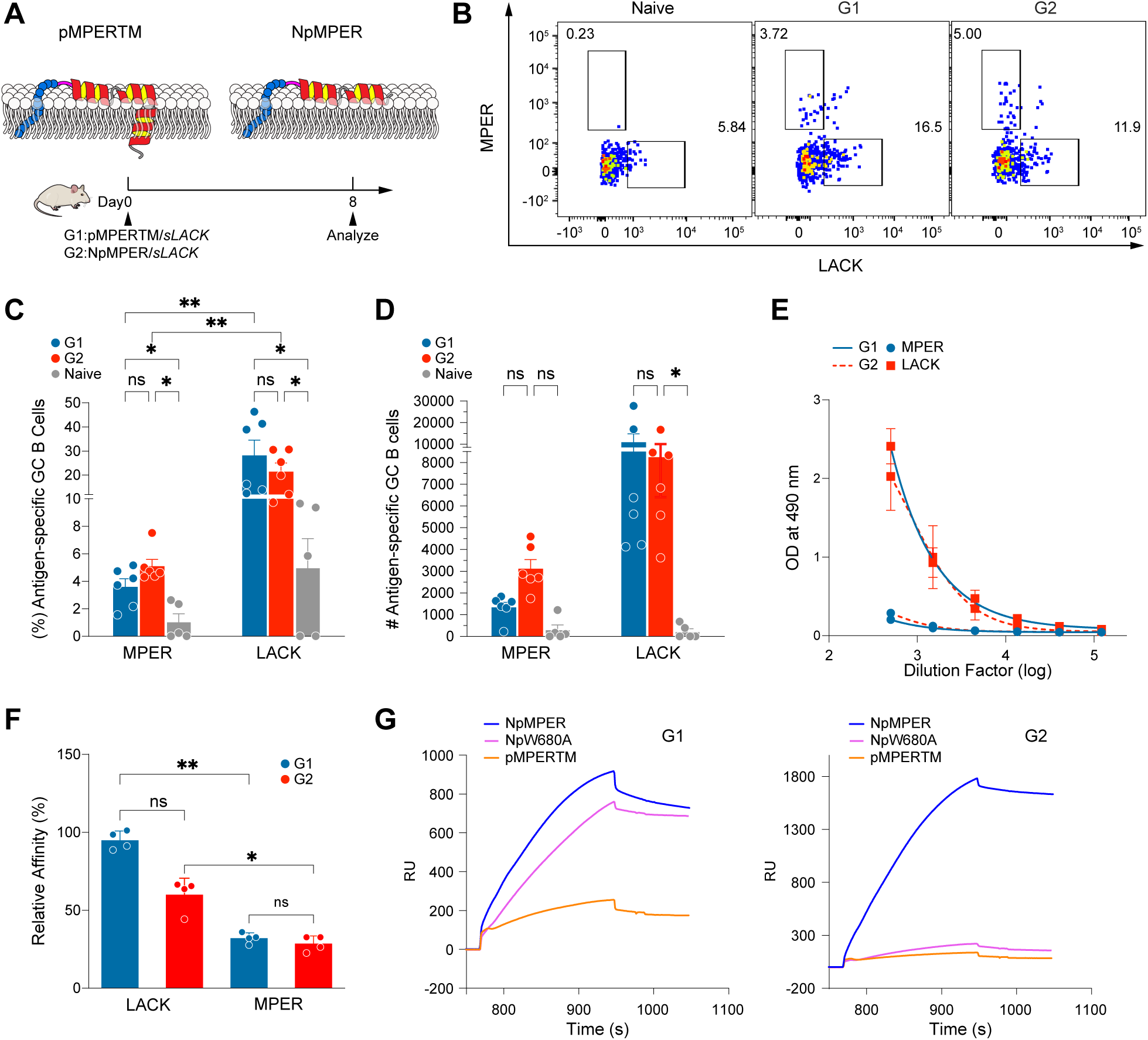
Dominant immune responses directed to the LACK helper peptide elicited by MPER/liposome vaccines. A) Schematic representation of MPER immunogens arrayed on the surface of liposome vaccine. B) Representative flow cytometry staining of MPER (B220^+^IgD^-^CD38^-^GL7^+^MPER-PE^+^) and LACK (B220+IgD^-^CD38^-^GL7^+^LACK-FITC^+^) specific GC B cells. Gating strategy is shown in Supplementary Fig. 1B. C-D) Frequency (C) and total number (D) of MPER- and LACK-specific GC B cells in the spleen assessed on day 8 after primary immunization in vaccinated mice (n = 6). Naïve mice (n = 5) were included as negative controls (also see Supplementary Fig. 1B-C). Data is represented as mean ± SEM (n=6) and results are representative of two independent experiments. Statistical differences were assessed by two-way ANOVA (or mixed model) with Tukey’s multiple comparison test *p < 0.05, **p < 0.01. E) MPER- and LACK-specific IgG titers from the immune sera collected on day 30 after immunization with pMPERTM or NpMPER formulated with sLACK helper peptides. F) Relative affinities of MPER- and LACK-specific serum IgGs collected 30 days after homologous boost immunization assessed by ELISA against NpMPER/liposome and pLACK/liposome, respectively. Calculation of the % relative affinity is detailed in Methods section. Statistical differences were assessed by Kruskal-Wallis test with Dunn’s comparison test denoted by p values: *p < 0.05, **p < 0.01. G) Antigenicity of the purified polyclonal IgG antibodies elicited by G1 and G2 immunogens. Surface plasmon resonance (SPR) sensograms depict the relative binding reactivity of the elicited antibodies for NpMPER/liposome, pMPERTM/liposome and NpW680A/liposome. N-terminally palmitoylated irrelevant NP366/liposome (Influenza A nucleoprotein (NP) peptide epitope) was used as a negative control for subtraction. Data shown is representative of three independent experiments.

Given that the sLACK contains potential B cell epitopes that are recognized by the immune system and play a key role in activating B cell responses, the antigen-specific germinal center (GC) responses were analyzed by flow cytometry using fluorescently conjugated NpMPER/liposome-PE or palmitoylated-LACK (pLACK)/liposome-FITC (lipid-fluorophore detailed in methods section (Fig. 1B and Supplementary Fig. 1B). The percentage and absolute number of MPER-specific GC B cells were greater following immunization with NpMPER/liposomes than that with pMPERTM/liposome (3.59% vs 5.09 % and 1329 vs 3111, respectively) (Fig. 1C and 1D). The total population of GC B cells was also larger for G2 than that for G1 (Supplementary Fig. 1C).

Surprisingly, LACK-specific B cells were recruited to the GC with increased frequency relative to MPER-specific B cells reaching 28.1% on day 8 for G1 and 21.3% for G2 and accompanied by a greater absolute number of cells as well (Fig. 1C and 1D). Consequently, the frequency of LACK-specific GC B cells coincided with serum antibody immunodominance directed against LACK (Fig. 1E) at 30 days after priming. Further, the extent of affinity maturation of the elicited antigen-specific serum IgG antibodies was assessed by ELISA assay based on the difference in binding of antibodies to the peptides at the low density (1:500 peptide:lipid ratio) versus at the high density (1:50 ratio) as previously established ^62^ (detailed in Methods). A considerable advantage in affinity maturation of the LACK-specific B cells compared to the MPER-specific B cells was evident. The relative affinity maturation of G1-LACK (94.8 %) and G2-LACK (59.9) were both high (Fig. 1F). On the other hand, affinity maturation against the MPER was low (°30%) and there was no significant difference between MPER-specific B cells between G1 and G2 (Fig. 1F). Importantly, the affinity maturation was significantly better against LACK than against the MPER. The immunodominant antibody response was maintained regardless of differences in MPER conjugation strategy and sequence used. Therefore, the results suggest that intrinsic affinity of BCRs for the LACK is higher than that of MPER immunogens.

We next compared the antigenicity of the purified polyclonal IgG antibodies elicited by NpMPER/liposomes vs pMPERTM/liposomes. While the binding of purified polyclonal IgG antibodies from G2 to NpMPER immunogen is relatively high, the antibody binding for W680A mutant MPER (NpW680A) and pMPERTM was abolished as revealed by surface plasmon resonance (SPR) analysis (Fig. 1G, right panel). The result confirms that the dominant antibody response in G2 is directed towards the C-terminal W680 residue and the NH_2_ chemical group as previously determined and noted above ^59^. On the other hand, similar binding reactivity for NpMPER and NpW680A was obtained by the purified polyclonal IgG antibodies elicited by pMPERTM/liposome (Fig. 1G, left panel). The results suggest that the pMPERTM can serve as a better priming immunogen, given that the pMPERTM-induced memory B cells imprint their epitope specificity free of NH_2_ or CH_3_CO group at the C-or N-terminus of the peptide immunogens.

### Heterologous boosting with successive changes in MPER and helper epitope peptides promote the expansion of long-lived plasma cells augmenting the MPER-specific serum antibody response

While the potential of pMPERTM as a prime immunogen is valid, the magnitude and affinity of pMPERTM-elicited serum antibodies are suboptimal. Given the LACK peptide immunodominance in generating humoral responses, we first tested whether heterologous boosting with different MPER immunogens and/or various helper epitope peptides could refocus immune responses away from the LACK. As shown in Fig. 2A, control group (G1) was immunized three times with pMPERTM/sLACK, whereas mice primed with pMPERTM/sLACK were boosted first with pMPERTM/soluble HIV30 (sHIV30) helper peptides and subsequently with pMPERTM/soluble OVA _323–339_ (sOVA) helper peptides in the place of sLACK in G2. Since a relative binding affinity of pMPERTM-elicited antibodies are much higher for NpMPER and NpW680A (Fig. 1G), the NpMPER and the NpW680A immunogens were also selected as a heterologous booster immunogen along with homologous sLACK (G3). In addition, the effect of heterologous B cell and CD4 T cell antigen boosting first with NpMPER/sHIV30, followed by NpW680A/sOVA was also compared (G4). Although the NpMPER, pMPERTM, and NpW680A immunogens are nearly identical in sequence (differing only by a single W680A mutation in NpW680A), they elicited distinct polyclonal antibody responses, as shown in Supplementary Fig. 2A. This difference in epitope specificity justifies our use of the term heterologous immunization.

**Fig. 2.**
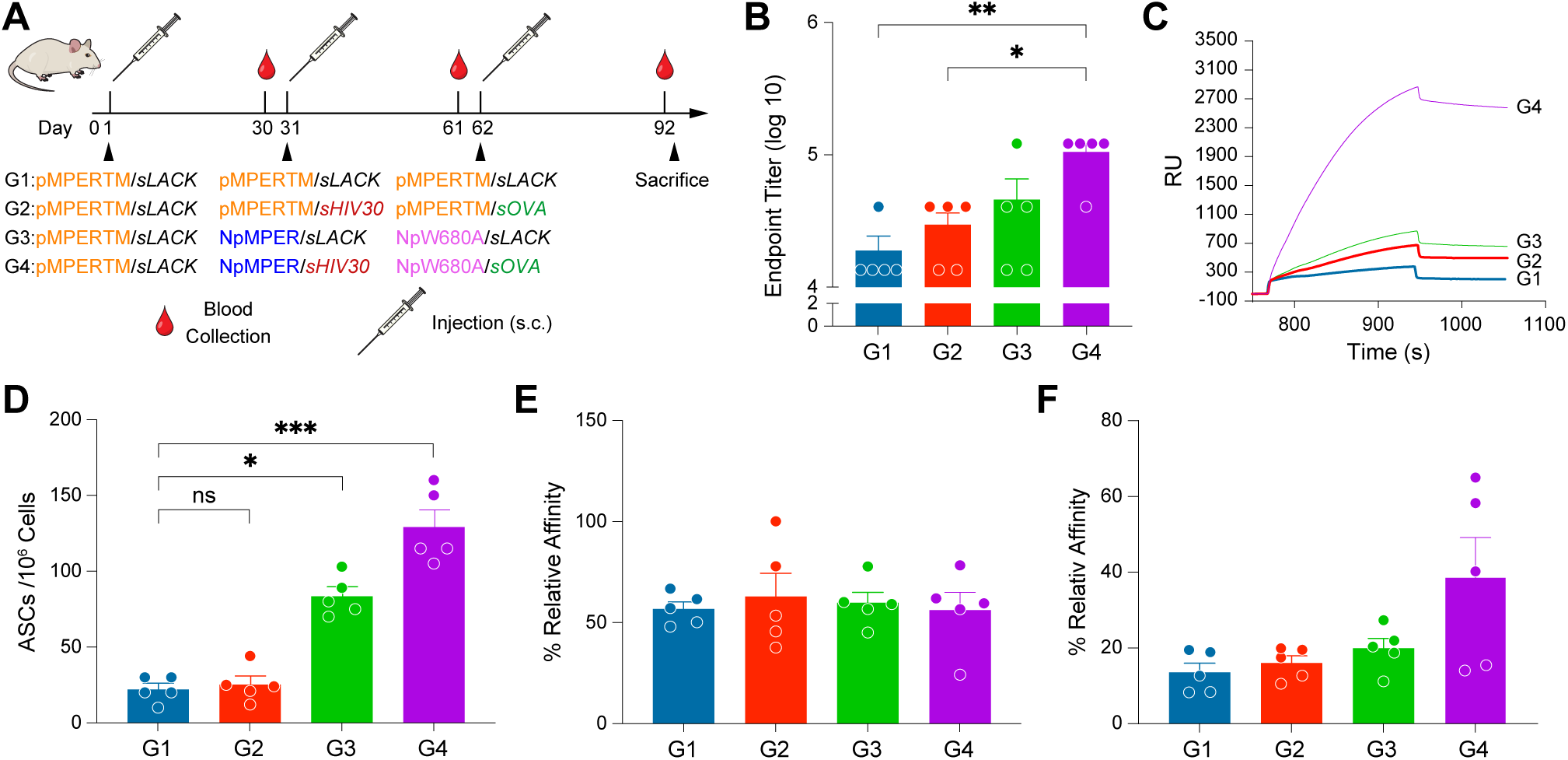
Comparison of MPER-specific immune responses elicited by various heterologous immunization regimens. A) Schematics of immunization schedule and heterologous immunization regimens. Balb/c mice were immunized subcutaneously with various MPER/liposome vaccines formulated with helper peptides and adjuvant at the specified time points. B) MPER-specific IgG endpoint titers (log10) from ELISA of immune sera against NpMPER/liposome collected 30 days after the third immunization (n=5). Data are representative of two independent experiments. Error bars represent mean ± SEM. Statistical differences were assessed by Kruskal-Wallis test with Dunn’s comparison test denoted by p values *p < 0.05, **p < 0.01. C) Relative binding activity of the purified polyclonal IgG antibodies elicited by each group for high density MPER/liposome (1:50) assessed by SPR analysis. The NpMPER/liposome was captured on L1 chip surface and sensorgrams illustrate the specific binding signal, measured in response units (RUs), on the y-axis as a function of time during the association and dissociation phases on the x-axis for each antibody at the concentration of 30ug/ml. Data shown are representative of three independent experiments, with non-specific antibody binding to NP366/liposome subtracted. D) Frequency of MPER-specific ASCs in bone marrow 30 days after the final immunization with each group indicated in A. Values represent background-subtracted data, with baseline ASC frequencies from naïve mice subtracted from each experimental group. Numbers of MPER-binding ASCs were determined by ELISPOT assay using high density NpMPER/liposome (1:50) as the capture antigen. Data is represented as mean ± SEM from one of three independent experiments (n=5). Statistically significant differences between different groups were determined by Kruskal-Wallis test with Dunn’s comparison test denoted by p values: *p < 0.05, ***p < 0.001 E) % relative affinities of antibodies secreted from ASCs measured by ELISPOT assay. The relative affinities were calculated by determining the high density MPER/Liposome (1:50) and low density (1:500) MPER/liposome binding ASCs from bone marrow cells. Data is represented as mean ± SEM from one of the three independent experiments (n=5). No statistically significant differences were observed. F) % relative affinities of MPER-specific IgGs in serum collected 30 days after the third immunization were assessed by calculating the ratio of MPER/liposome binding at low density (1:500) to high density (1:50) using ELISA (detailed in Methods section). The data is presented as mean ± SEM from one of two independent experiments, each with n = 5. No statistically significant differences were observed.

Mice in each group were first primed with pMPERTM/sLACK followed by boosting twice with the indicated various MPER and/or the helper peptide antigens (Fig. 2A). Immune sera collected 30 days after the final immunization were characterized to compare magnitude and quality of the elicited antibodies.

Overall, the MPER-directed serum titer was increased by ELISA measured against NpMPER/liposome antigen in all groups boosted with heterologous MPER and/or heterologous helper peptides compared to the control group boosted with homologous pMPERTM/sLACK (G1) (Fig. 2B and Supplementary Fig. 2B). However, significantly higher anti-MPER serum IgG titer (5.5-fold) was elicited when mice were boosted with both the heterologous MPER and the heterologous helper epitope peptides (G4) compared to heterologous MPER and homologous sLACK in G3 (2.4-fold) and the homologous pMPERTM and the heterologous helper epitope peptides in G2 (1.5-fold) (Fig. 2B). A relative binding affinity of purified polyclonal IgG antibodies for NpMPER/liposome assessed by SPR analysis also trended with the ELISA results and was most evident for G4 (Fig. 2C). While high-titer anti-LACK antibody responses were sustained following homologous boosting with sLACK in groups G1 and G3, these responses declined after heterologous boosting with sHIV30 and sOVA in groups G2 and G4 (Supplementary Fig. 2C) by ELISA using the purified polyclonal antibodies. In contrast, the antibody responses against HIV30 and OVA were generally weak.

As plasma cells (PCs) are a source of circulating serum antibody levels, we next tested the effect of each immunization regimen on the generation of MPER-specific long lived plasma cells (LLPC). Thirty days after the final immunization, bone marrow (BM) cells were isolated and cultured in wells coated with NpMPER/liposome to assess the frequency of antibody secreting cells (ASCs). Enzyme-linked ImmunoSpot (ELISPOT) analysis demonstrated that the frequency of ASCs (spots/million cells) in BM is considerably higher in G4 (mean=129) compared to G3 (83), G2 (25) and G1(22) (Fig. 2D). MPER-specific ASCs were not detected from naïve BM cells (data not shown). Therefore, the 5.8-fold increase in ASCs in BM for G4 correlates well with the 5-fold increase in serum antibody compared to that in the homologous immunization control group G1. Further, the analysis of antibody avidity for MPER/liposome within ASCs in BM estimated that the percentage of cells binding both NpMPER/liposomes at 1:500 and NpMPER/liposomes at 1:50 (peptide to lipid ratio, mol: mol) reached nearly 50-55% in all groups (Fig. 2E). We also assessed affinity maturation of serum antibodies, considering that short-lived plasma cells contribute to this pool, and that some plasma cells—potentially derived from memory B cells and residing in tissues—may also play a role in maintaining circulating antibody levels. Although the affinity/avidity of anti-MPER serum antibody was enhanced by about 2-fold by boosting with the heterologous MPER and the heterologous helper peptides (G4) compared to the homologous immunization (G1), the affinity maturation rate of anti-MPER serum antibodies achieved 38% in total with G4 (Fig. 2F), suggesting that a majority of ASCs may have produced low affinity antibodies.

### Epitope mapping analysis suggests clonal selection and expansion of MPER-specific antibodies by heterologous boosts with successive changes in MPER and helper epitope peptides

We next assessed the qualitative difference in the polyclonal antibody responses elicited by each immunization group. The resulting purified serum IgGs collected at 30 days after the final boost were first investigated for MPER-N (a.a 662-676) vs MPER-C segment (a.a 667-683) specific antibody responses by SPR. The relative binding reactivity of the purified polyclonal IgGs for MPER-N/liposomes was comparable to that for MPER-C/liposomes when mice were boosted with homologous pMPERTM/sLACK (G1) or when mice were boosted with NpMPER/sLACK and NpW680A/sLACK (G3) (Fig. 3A). Boosting with pMPERTM/sHIV30 and pMPERTM/sOVA (G2) or with NpMPER/sHIV30 and NpW680A/sOVA immunogens (G4) induced a dominant antibody response directed to MPER-N domain (Fig. 3A). To quantify MPER-N or MPER-C specific antibodies produced by ASCs in BM, BM cells were isolated for ELISPOT analysis 30 days after the final immunization. The frequency of MPER-N-specific ASCs was highest in G4 (84) followed by G3 (39)> G2 (23)> G1(12) spots/million cells, whereas the frequency of MPER-C specific ASCs was highest in G3 (40) >G4 (19)>G1 (11)>G2 (10) spots/million cells (Fig. 3B). Although the magnitude of total ASCs was influenced by the context of both the MPER and the helper peptide immunogens, the recruitment of MPER-C-specific B cells appears to be influenced by the helper peptide function (Fig. 3C).

**Fig. 3.**
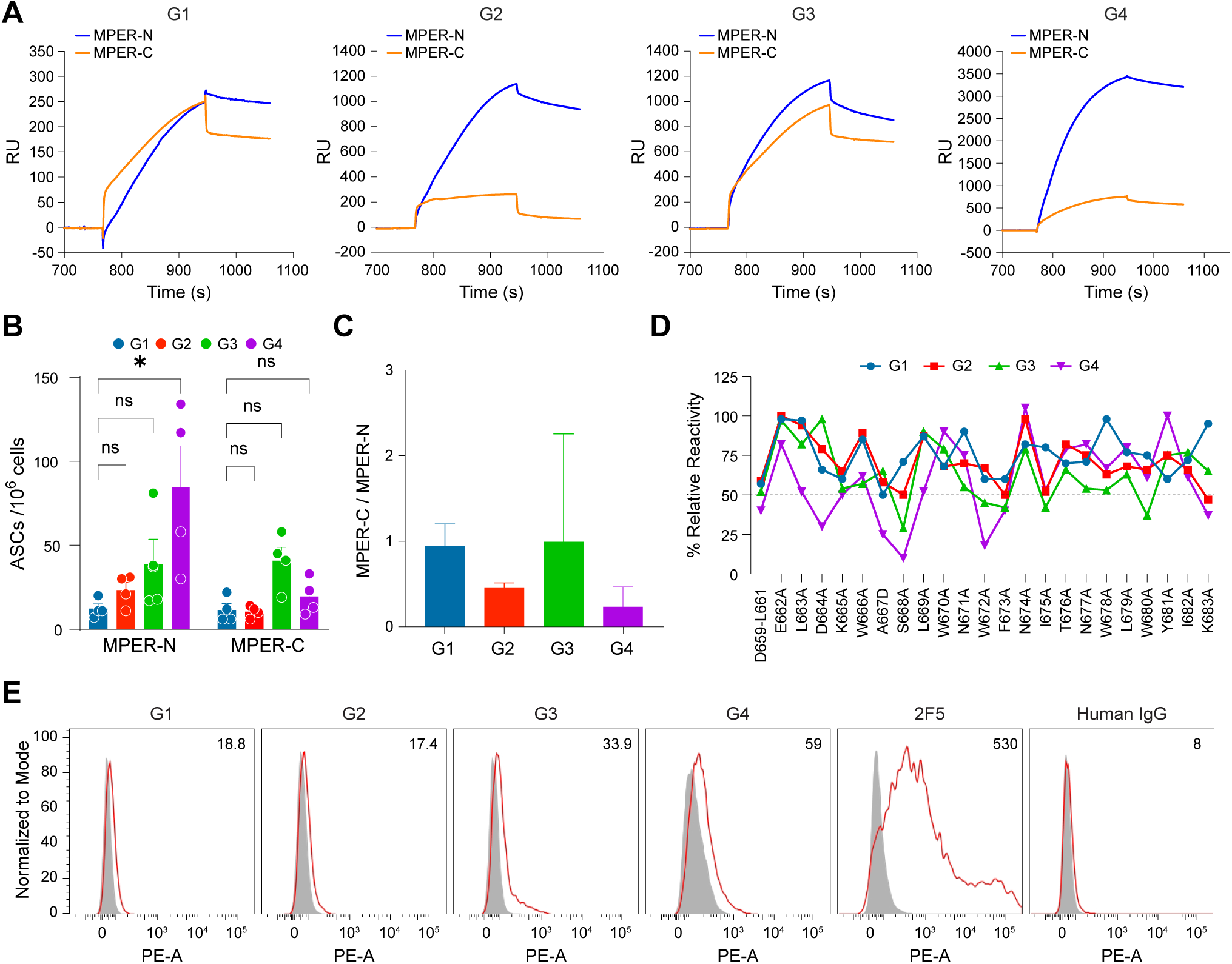
Characteristics of MPER-specific antibodies elicited by various heterologous immunizations. A) Qualitative binding analysis of MPER-N and MPER-C domain-specific antibodies elicited by the indicated groups in Fig. 2A. SPR sensograms are shown for the purified polyclonal antibody binding from each group to MPER-N/liposome and MPER-C/liposome. The antibodies were tested at a concentration of 30-40 µg/ml by injection over L1 chip-bound MPER/liposomes at a flow rate of 20 µl/min for 3 min. MPER-N is shown in dark blue, while MPER-C is in tan. Data shown are representative of three independent experiments, with non-specific antibody binding to NP366/liposome subtracted. B) Frequencies of MPER-N and MPER-C domain-specific ASCs in bone marrow measured by ELISPOT. Bar graph represents the quantification of ASCs calculated from 10^6^ bone marrow cells from different groups as indicated in Fig. 2A. Statistically significant differences between the various groups were determined by two-way ANOVA with Tukey’s multiple comparison test denoted by p values: *p < 0.05 C) The ratio of MPER-C vs MPER-N-specific ASCs based on the frequencies of ASCs estimated in Fig. 2B. Statistically significant differences between different groups were determined by Kruskal-Wallis test with Dunn’s comparison denoted by p values: **p < 0.01. Data in panels B and C shown are a representative of two independent experiments with individual animals per group (n=4). D) Epitope mapping analysis by BIAcore of the purified polyclonal IgG antibodies from each group by use of liposome-bound serial single alanine MPER mutants. The y axis shows the percent relative binding activities of antibodies against each single-residue mutant compared to that of wild-type MPER (100%). The % relative binding activities of antibodies against D659-L661 was calculated based on the difference in binding response units for wild type D659-L661 MPER (a.a 659-682) and for wild type MPER (a.a 662-682). E) ADA gp145 binding by the various vaccine-elicited antibodies along with a known bnAb 2F5 as a positive control by flow cytometry. Anti-human polyclonal IgG served as negative control. Background staining of Abs to untransfected cells is shown in gray and MFI values are indicated after subtraction of the antibody binding from each group to untransfected 293T cells from each group. A representative set of histograms of n=2 biologically independent experiments is shown.

To delineate the key epitope MPER residues recognized by the purified polyclonal IgG antibodies from each group, a series of MPER mutants harboring single alanine substitutions were tested for the polyclonal antibody binding. A dominant contribution of MPER residues defining characteristics of antibody specificity appeared lacking when pMPERTM/sLACK-primed mice were boosted either with the homologous pMPERTM/sLACK antigens (G1) or with the homologous pMPERTM and the heterologous helper peptides (G2), as shown in Fig. 3D. On the other hand, polyclonal antibody binding to MPER mediated by residues such as S668, F673, I675, W680 was apparent with each residue binding contribution ranging from 58-71% when the primed mice were boosted with the heterologous MPER and the homologous sLACK peptides (G3). Interestingly, the boost regimen with the heterologous MPERs and the heterologous helper epitope peptides (G4) elicited polyclonal antibody epitope specificity defined distinctively by a dominant contribution of residues D664, A667, S668 and W672 (Fig. 3D), consistent with the frequency of ASCs producing antibodies directed to MPER-N domain exhibited in the serum antibodies (Fig. 3A and B). Furthermore, the effect of the different boosting regimens on the function of antibody was also evident as the heterologous MPER immunogens (G3 and G4) compared to the homologous MPER immunogens (G1 and G2) induced improved antibody binding to ADA gp145 with its MPER sequence matching the NpMPER and pMPERTM immunogen sequence expressed on the surface of 293T cells. Specifically, the heterologous G4 and G3 regimens resulted in a 3.1-fold and 1.8-fold increases, respectively, compared to the homologous G1 regimen MFI values (Fig. 3E).

### Secondary GC response *per se* is not correlated with the outcome of anti-MPER antibody responses

Since the immunization regimen for G4 induced MPER-specific antibody responses greater in quality and quantity compared to the other groups, we next sought to understand the underlying cellular mechanisms for the superiority. Follicular T helper (Tfh) cells play an essential role in regulating GC reaction and consequently generating robust serum antibody responses ^63^. Given that the difference in the immunization regimens between G3 and G4 (Fig. 2A) is the homologous vs. the heterologous helper epitope peptides formulated in the booster vaccines, we investigated the magnitude of the secondary GC responses mediated by the different CD4 T cell helper peptides. The GC responses were measured by flow cytometry from spleens of mice 12 days post-secondary immunization at day 43. The NpMPER/sLACK boosting (G3) led to larger total Tfh (PD-1hi/CXCR5^+^) cell frequency and numbers as well as GC B cell responses when compared to G4 boosted with the NpMPER/sHIV30 (Fig. 4A, 4B and Supplementary Fig. 3). GC B cells were also stained with PE-labeled MPER/liposome and FITC-labeled LACK/liposome or PE-labeled MPER/liposome and FITC-labeled HIV30/liposome to assess each antigen-specific GC B cells, respectively. The frequency of MPER-specific GC B cells was slightly higher in G4 than that in G3, although not statistically significant (Fig. 4C). However, the MPER-specific total GC B cell number was higher in G3 compared to that in G4 due to smaller total number of GC B cells. On the other hand, significantly more LACK-specific GC B cells than the MPER-specific GC B cells were recruited to GC reactions when mice in G3 were boosted with the sLACK (5.2% vs 1.3 %), whereas sHIV30-specific GC B cells were slightly higher than MPER-specific GC B cells in G4 boosted with the sHIV30 helper peptides (3.98 vs 1.85%) (Fig. 4C).

**Fig. 4.**
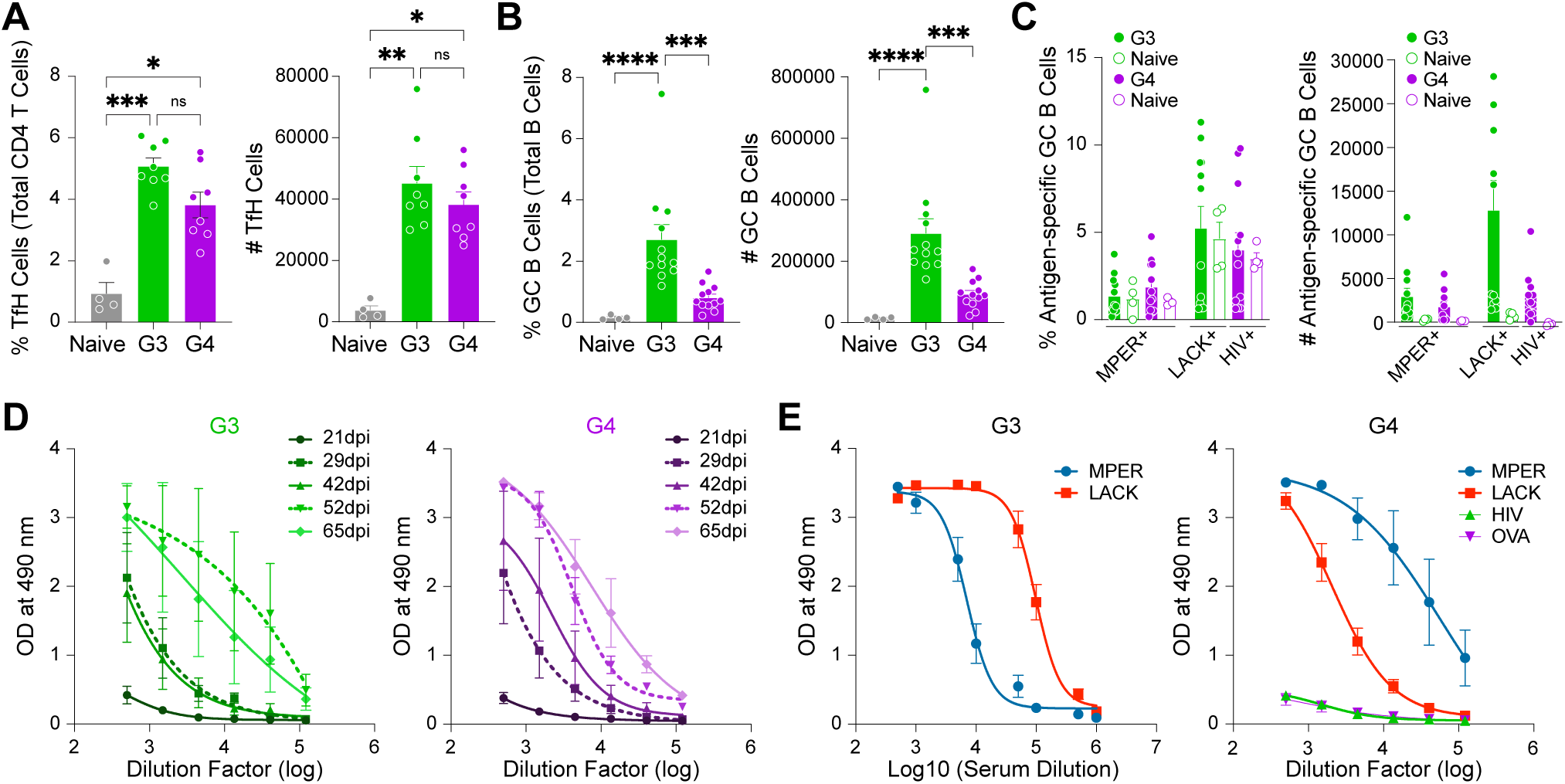
Secondary GC responses following boost with MPER immunogen formulated with homologous vs heterologous helper epitope peptides. A-C) Frequencies and numbers of Tfh (CD4^+^CD44^+^CD62L^-^PD1^+^CXCR5^+^) cells following booster immunization (A) (also see Supplementary Fig. 3 for gating strategy), GC B cells (B220^+^IgD^-^CD38^-^GL7^+^) (B), and antigen-specific GC B cells (C). MPER- (B220^+^IgD^-^CD38^-^ GL7^+^MPER-PE^+^) and LACK- (B220^+^IgD^-^CD38^-^GL7^+^LACK-FITC^+^) specific GC B cells or MPER- (B220^+^IgD^-^CD38^-^GL7^+^MPER-PE^+^) and HIV30- (B220^+^IgD^-^CD38^-^GL7^+^HIV-FITC^+^) specific GC B cells quantified by FACS and presented in bar graphs as percent and total cell number. For panels A-C, Balb/c mice primed with pMPERTM/sLACK were boosted with NpMPER/sLACK (G3) or NpMPER/sHIV30 (G4). The Tfh and GC responses induced by G3 and G4 were assessed by flow cytometry from spleen on day 12 after the boost. Symbols represent individual animals, and error bars indicate mean ± SEM. The data are cumulative from two independent experiments. Unimmunized mice were used as controls. Statistical significance for the panel A and B was determined using Kruskal-Wallis test with Dunn’s comparison test with p-values indicated as, *p < 0.05, **p < 0.01, ***p < 0.001, ****p < 0.0001. D) MPER-specific IgG titers from immune sera collected at the indicated timepoints. The broken lines indicate immune sera collected on day 29 and day 52 after primary immunization, respectively. The data are represented as binding curves, with log serial dilutions of serum on the x-axis and absorbance values on the y-axis. E) ELISA to measure serum IgG titers for MPER/liposome, LACK/liposome, HIV30/liposome and OVA/liposome. The immune sera were collected 30 days after the third immunization and LACK- vs MPER- specific antibody titers induced by G3 regimen (left) are compared with MPER-, LACK-, HIV30- and OVA-specific antibody titers induced by G4 regimen (right). All peptides were N-terminally palmitoylated and arrayed on the surface of liposome for ELISA assay.

Given that GC response is closely related to the future antibody response, recall responses were measured subsequently for anti-MPER antibody titers by ELISA. Priming of G3 and G4 with the same pMPERTM/sLACK elicited low antibody titers (Fig. 4D). Boosting with the homologous sLACK in G3 induced antibody memory response with anti-MPER titer rising until day 29 but plateauing by day 42. In contrast, the heterologous boosting with sHIV30 in G4 led to a continued increase in anti-MPER titer from day 29 to day 42 (Fig. 4D). The same trend was observed when mice in G3 and G4 were boosted the second time with W680A/sLACK and W680A/sOVA (Fig. 4D). Thus, while the sLACK exerted stronger CD4 T cell help over sHIV30 to induce greater secondary GC responses, the greater anti-MPER antibody response was linked to boosting with heterologous sHIV30 than with homologous sLACK despite a smaller overall GC response. The dominant LACK-specific secondary GC response boosted further the anti-LACK antibody titer following two homologous boosts with the sLACK in G3 (Fig, 4E, left panel). In contrast, boosting with the sHIV30 and the sOVA peptides induced weak anti-HIV30 and anti-OVA antibody responses compared to the anti-MPER antibody response in G4 (Fig. 4E, right panel). The difference may be attributed to the lack of recall antibody responses mediated by HIV30- and OVA-specific memory B cells. Alternatively, or in addition, it may suggest that the intrinsic B cell affinity for the HIV30 and the OVA B cell epitopes stimulated against the helper peptides may be much lower than that for the LACK.

### MPER-specific antibody responses are modulated by the immunogenicity of B cell epitopes for the helper peptides

We next examined the capacity of sLACK, sHIV30 and sOVA CD4 T cell help to recruit MPER-specific GC B cells from spleens 12 days after mice were primed with NpMPER formulated with sLACK, sHIV30 or sOVA, respectively, (denoted as sG1, sG2 and sG3). When compared to sHIV30 and sOVA, sLACK resulted in a statistically significant increase in the percentage of Tfh cells (Fig. 5A). The corresponding GC B cell response showed less pronounced differences between the sLACK group and the sHIV30 and sOVA groups (Fig. 5B). In terms of antigen specificity within the GC B cell population, the LACK-specific response was clearly more dominant than the MPER-specific response (5.4 % vs 2.4 %), unlike the more comparable MPER- vs. HIV30-specific (1.7 % vs 1.9 %) and MPER- vs OVA-specific responses (2.9 % vs 3.4%) (Fig. 5C, left panel). The dominant LACK-specific GC response, also reflected in the greatest number of antigen-specific GC B cells trend (Fig. 5C, right panel) was consistent with greater anti-LACK antibody titer compared to anti-MPER antibody titer (Fig. 5D, panel sG1). On the other hand, although sHIV30- and sOVA-specific GC B cells were slightly higher than the MPER-specific GC B cells, subsequent anti-HIV30 and anti-OVA antibody responses were much weaker than anti-MPER antibody responses (Fig. 5D, panels sG2 and sG3).

**Fig. 5.**
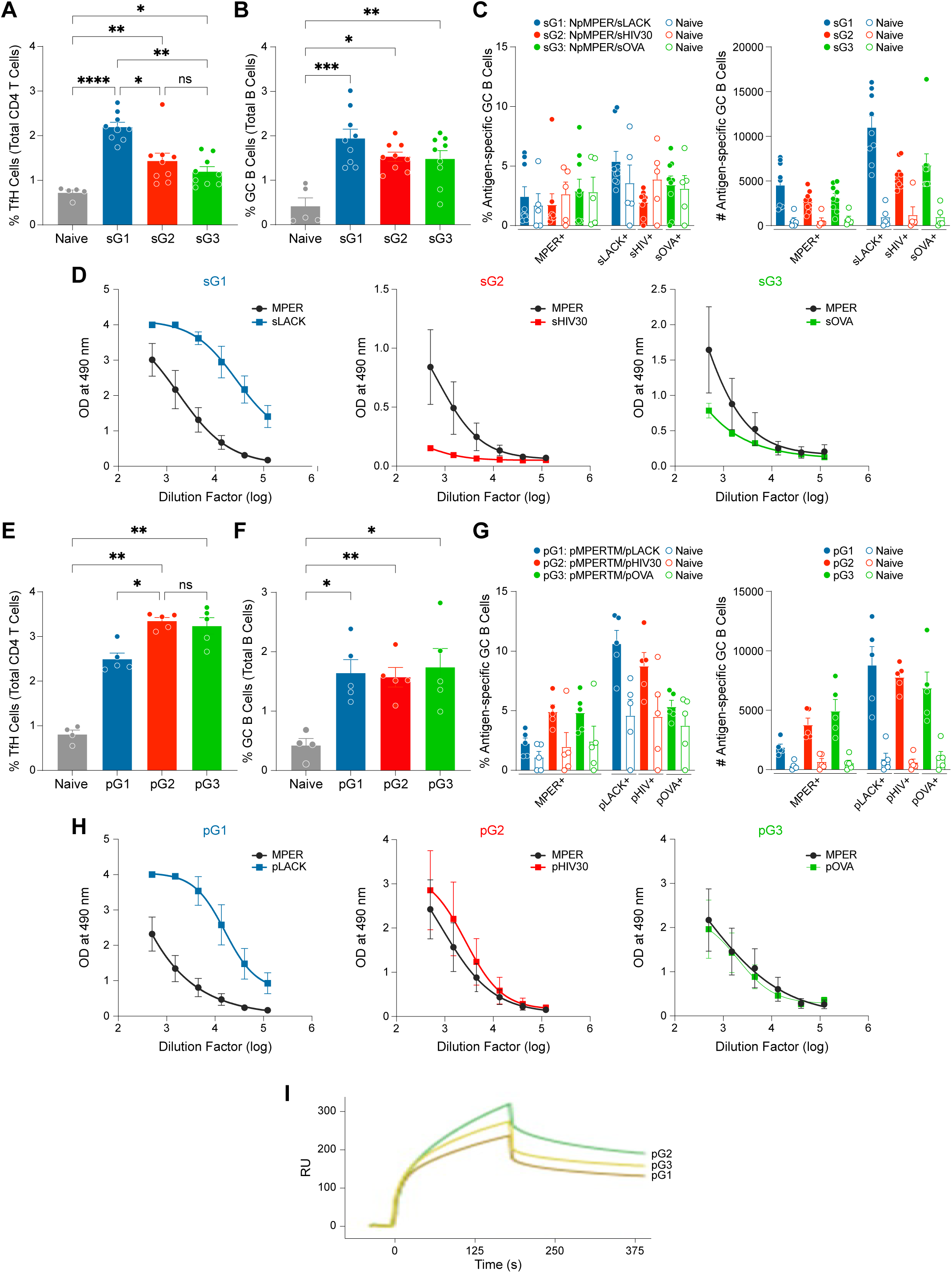
Effects of various helper epitope peptide formulations on the efficacy of CD4 T cell help and antigen-specific antibody responses. A-C) Flow cytometric analysis and quantification of the percentage of Tfh cells (A), GC B cells (B) and percentage and total cell number of antigen-specific GC B cells (C) in the spleens of Balb/c mice on day 12 post primary immunization. Statistical significance was determined using Kruskal-Wallis test with Dunn’s comparison test, with p-values indicated as, *p < 0.05, **p < 0.01, ***p < 0.001, ****p < 0.0001. Mice were immunized with NpMPER/liposome with sLACK (sG1), sHIV30 (sG2) or sOVA helper peptides (sG3) encapsulated inside liposome vaccine, respectively. Naïve mice (n=5), served as controls. Error bars represent mean ± SEM from n = 9 animals, based on one of the two independent experiments. (D) Comparison of antigen-specific serum IgG titers measured by ELISA after homologous boosting. MPER-specific IgG titers from immune sera of mice in each group collected 10 days after a single homologous boost was compared with anti-LACK (sG1), anti-HIV30 (sG2) or anti-OVA antibody titers (sG3). (E-G) GC responses induced by co-delivery of helper peptides pLACK, pHIV30 or pOVA with NpMPER/liposome at primary response. Frequency of Tfh cells (E), frequency of GC B cells (F) and frequency and number of antigen-specific GC B cells (G) in the spleen of Balb/c mice on day 12 after primary immunization with NpMPER/pLACK (pG1), NpMPER/pHIV30 (pG2) and NpMPER/pOVA (pG3). Responses in naïve mice (n = 4) were assessed as controls. Data shown are representative of two independent experiments (n=5). Statistical significance was determined using Kruskal-Wallis test with Dunn’s comparison test, with p-values indicated as, *p < 0.05, **p < 0.01. (H) Immunogenicity of three different helper peptide antigens relative to the MPER in indicated formulations after homologous boosting. MPER-specific IgG titers by ELISA are compared across the three groups, each corresponding to the specific CD4 T cell helper peptide pLACK (pG1), pHIV30 (pG2), and pOVA (pG3). The data are presented as binding curves, with log serial dilutions of serum on the x-axis and absorbance values on the y-axis. I) A relative binding affinity of the purified polyclonal IgG antibodies for MPER/liposome. The MPER/liposomes at low density (peptide to lipid ratio of 1 :500) were captured on the L1 chip surface and the polyclonal IgG antibodies elicited by each group were passed over the chip surface for 3 mins. The polyclonal IgG antibodies from each group were purified from immune sera collected 10 days after boost. Data shown are representative of three independent experiments, with non-specific antibody binding to NP366/liposome subtracted.

The magnitude of CD4 T cell help is directly correlated with antigen amount ^64^. Since CD4 T cell helper peptide encapsulation into liposome can be greatly influenced by the peptide sequence, we speculate that differences in the Tfh cell responses induced by the three different helper peptides could be attributed to the helper peptide encapsulation efficiency. To that end, we assessed the effect of liposome vaccine formulation on the recruitment of antigen-specific GC B cells. Three different groups of mice were primed with liposome vaccines arraying NpMPER and N-terminally palmitoylated CD4 T cell helper peptides (pLACK, pHIV30 or pOVA, and denoted as pG1, pG2 and pG3), respectively. In contrast to the sHIV30, the frequency of Tfh cells from spleen of mice 14 days after the prime with NpMPER/pHIV30 was slightly higher than that induced by pLACK and pOVA (Fig. 5E). GC frequency was comparable to each other (Fig. 5F), suggesting that the delivery of HIV30 and OVA peptides covalently anchored in a liposome augmented the magnitude of GC responses compared to the delivery of sHIV30 and sOVA encapsulated into liposome, respectively (Fig. 5F, pG2 and pG3 vs. Fig. 5B, sG2 and sG3). pHIV30 and pOVA delivery also increased the frequency of MPER-(4.9% for pHIV30 vs 1.7% for sHIV30 and 4.8% for pOVA vs 2.9% for sOVA), HIV30-(8.8% for pHIV30 vs 1.9% for sHIV30) and OVA-specific (5.3% for pOVA vs 3.4% for sOVA) GC B cells, respectively compared to sHIV30 and sOVA delivery (Fig. 5G vs. 5C). In contrast, while pLACK increased the frequency of LACK-specific GC B cells 2-fold compared to sLACK (10.6% vs 5.4%), the MPER-specific GC response remained similar (Fig. 5G vs. 5C). This disparity resulted in a more dominant LACK-specific GC B cell hierarchy after pLACK delivery (2.2% MPER vs. 10.6% LACK) compared to sLACK (2.4% MPER vs. 5.4% LACK). While the hierarchy of MPER- and OVA-specific GC responses remained similar, pHIV30 delivery was pronounced with greater HIV-30 specific GC B cells than MPER-specific GC B cells (8.8% vs 4.9%) compared to that induced by sHIV-30 delivery (1.7% vs 1.9%) (Fig. 5G and Fig. 5C).

To further characterize the quality of helper T cell responses elicited by the different peptides pG1, pG2 and pG3, we examined effector antigen-specific CD4⁺ T cell function ex vivo. Splenocytes were collected 10 days after a booster immunization and restimulated with their respective peptides for characterization. IFN-γ ELISPOT analysis of total splenocytes revealed similar frequencies of LACK and HIV30 antigen-specific IFN-γ–secreting cells and slightly higher frequency of OVA-specific IFN-γ–secreting cells, although statistically not significant (Supplementary Fig. 4A). Assessment of IFN-γ production among activated (CD44⁺) CD4⁺ T cells by FACS analysis showed comparable responses between LACK- and HIV30-specific cells, whereas OVA-specific responses were slightly lower (Supplementary Fig. 4B). Overall activated Tfh cell frequencies were also comparable across all groups (Supplementary Fig. 4C). Similarly, analysis of antigen-specific Tfh activation, identified by upregulation of CD25 and OX40, showed largely similar responses between LACK and HIV30, with a modest increase observed in the HIV30 and LACK groups compared to the OVA group (Supplementary Fig. 4D). The results imply that differential helper peptide availability, likely a consequence of the encapsulation efficiency of soluble helper peptides into liposome, accounts for the enhanced Tfh cell response observed with sLACK relative to sHIV30 and sOVA. Nevertheless, the augmented anti-MPER response induced by G4 regimen over the antibody response elicited by G3 regimen (Fig. 2A) seems to be uncorrelated with the stronger CD4 T cell help per se provided by boosting immunizations with sLACK compared to heterologous boosting with sHIV30 and sOVA.

Accordingly, we next tested the intrinsic B cell affinity for antigens in the context of MPER/liposome vaccines as a factor associated with the augmented anti-MPER responses. Subsequently, MPER-specific antibody responses were measured from immune sera collected 10 days post homologous booster immunization. Anti-MPER titer was similar in each immunization group (Fig. 5H). However, NpMPER/pLACK elicited dominant anti-LACK antibody titer, a trend similar to that elicited by sLACK, whereas anti-MPER antibody titer elicited by pLACK was slightly reduced compared to the antibody titer elicited by sLACK (Fig. 5H, pG1 panel vs Fig. 5D, sG1 panel). In contrast, anti-HIV30 and anti-OVA antibody titers were comparable to the anti-MPER antibody responses (Fig. 5H pG2 and pG3 panels). The augmented anti-HIV30 and anti-OVA antibody responses compared to that elicited by sHIV30 and sOVA (Fig. 5H vs Fig. 5D) are likely due to increased amounts of these antigens on the surface of liposome. Furthermore, the relative binding affinity of the purified polyclonal IgG antibodies for low density MPER/liposome (1: 500) assessed by SPR analysis suggest that the affinity of MPER-specific antibodies elicited by NpMPER/pLACK is slightly lower than that elicited by either NpMPER/pOVA or NpMPER/pHIV30 (Fig. 5I).

### Priming immunogen along with augmented CD4 T cell help shape the magnitude and the quality of the subsequent MPER-specific immune responses

In view of the augmented CD4 T cell help by pHIV30 compared to sHIV30, we next examined the impact of co-delivery of the membrane-anchored helper peptides and the MPER arrayed on the surface of liposome. Mice primed with pMPERTM/sLACK were either boosted first with NpMPER/sHIV30 and second with NpW680A/sOVA in G1 as a control or boosted with NpMPER/pHIV30 followed by NpW680A/pOVA in G2 (Fig. 6A). While all three helper epitopes exerted similar CD4 T cell helper function, LACK appears to be a more dominant B cell antigen compared to HIV30 and OVA (Fig. 5). Therefore, we also examined the impact of pMPERTM/pHIV30 priming in comparison with pMPERTM/sLACK. As indicated in Fig. 6A, mice primed with pMPERTM/pHIV30 were followed by two booster immunizations with NpMPER/pOVA and NpW680A/sLACK.

**Fig. 6.**
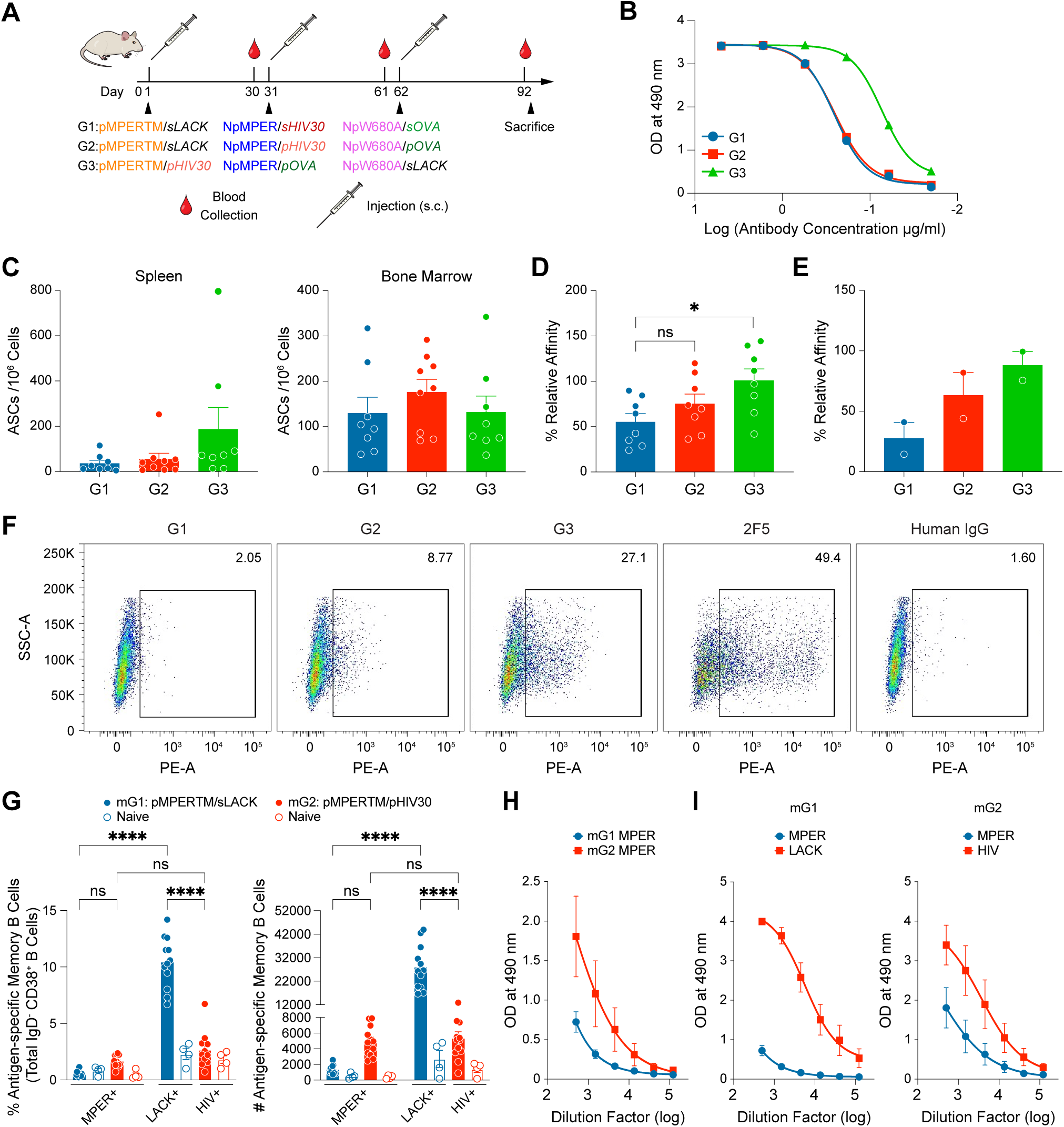
Effects of priming immunogen and augmented CD4 T cell help on the magnitude and quality of MPER-specific antibody responses. A) Schematic of heterologous immunization strategies. B) Anti-MPER polyclonal IgG titers determined against high density NpMPER/liposome (1:50) by ELISA. Immune sera were collected 30 days after the third immunization, combined from 5 mice for each group and purified using gamma-bind plus affinity column for the assay. Data shown is representative of two independent experiments. C) Frequency of MPER-specific ASCs in spleen and bone marrow. The MPER-specific ASCs from spleen (left) and bone marrow cells (right) at 30 days after the third immunization were determined from 8 mice by ELISPOT assay. Quantification of ASCs calculated from 10^6^ cells from each group is indicated. Data are background-subtracted using ASC frequencies from naïve mice as baseline controls. Error bars indicate mean ± SEM. D) % relative affinities of MPER/liposome binding IgG ASCs in bone marrow as assessed by ELISPOT assay. The frequencies of low-density MPER/liposome (1:500) and high density MPER/liposome (1:50) binding IgG ASCs were determined with the ratios of low density MPER to high-density MPER binding plotted as % relative affinity. Higher percentage is indicative of greater affinity antibodies. Data is represented as mean ± SEM. Statistically significant differences between different groups were determined using Kruskal-Wallis test with Dunn’s comparison test denoted by p values: *p < 0.05. E) % relative affinities of MPER-specific purified polyclonal IgG antibodies measured by ELISA against low density (1:500) vs. high density (1:50 peptide to lipid ratio) MPER/liposome as explained in details in Methods section. The polyclonal antibodies were purified from immune sera combined from 5 mice for each group collected on day 30 after the third immunization. No statistically significant differences were observed. F) Vaccine-elicited antibody binding to ADA gp145 expressed on the surface of 293T cells by flow cytometry. 293T cells were transfected to express ADA gp145 with MPER sequence mutated to the MPER immunogen (HxB2 MPER sequence), and binding was measured using anti-mouse IgG-PE. Flow cytometry plots show side scatter area (SSC-A) on the Y-axis and PE fluorescence (ADA gp145 binding) on the X-axis. The data are presented as a representative example from one of two independent experiments. G) Frequency and total cell number of MPER-specific memory B cells. Balb/c mice were immunized with pMPERTM/sLACK liposome (mG1) or pMPERTM/pHIV30 liposome (mG2), and the MPER-specific IgG memory B cells from each group was assessed in spleen on day 30 post-primary immunization. Data shown are represented as the aggregate of two independent experiments (n=12). Statistically significant differences between the groups were determined by two-way ANOVA (or mixed model) with Tukey’s multiple comparison test denoted by p values: ****p < 0.0001. H) Difference in quantity of anti-MPER IgG responses elicited by mG1 and mG2. Immune sera were collected 10 days after the homologous booster immunization. I) Comparison of MPER and helper peptide-specific antibody titers elicited by pMPERTM/sLACK (mG1) and by pMPERTM/pHIV30 (mG2). Data in H-I are represented as one of the two independent experiments with n=6.

Anti-MPER antibody titers, measured by ELISA using purified polyclonal IgGs collected 30 days after the final immunization, were comparable between groups G1 and G2. In contrast, group G3 showed a modest increase in anti-MPER titers relative to G1 and G2 (Fig. 6B). Further ELISPOT analysis demonstrated that the frequency of ASCs in BM trended slightly higher in G2 (176) compared to ASCs induced by G1 (129) and G3 (132) spots /million cells, although not statistically significant. On the other hand, increased numbers of ASCs resided in the spleen from mice in G3 (187) compared to that in G1 (36) and G2(55) (Fig. 6C). Additionally, augmented affinity maturation of MPER-specific antibodies produced from ASCs in BM elicited by G3 was pronounced compared to that elicited by G1 and G2 (Fig. 6D). Likewise, the affinity of anti-MPER antibodies in plasma was also improved when mice were boosted with pHIV30 and pOVA compared to sHIV30 and sOVA helper peptides. The maximal effect on the induction of high affinity antibodies for the MPER was associated with either the priming immunogen per se and/or order of heterologous immunogens rather than NpMPER/pHIV30 and NpW680A/pOVA modules (Fig. 6E). In line with the rate of antibody affinity maturation among the three groups, 27% of 293T cells expressing ADA gp145 were stained by G3-elicited purified polyclonal IgG antibodies followed by 8% and 2 % staining by the G2 and G1 equivalents (Fig. 6F).

To directly compare the priming efficiency of CD4 T cell helper peptides, we assessed the frequency of Tfh cells induced by pMPERTM/sLACK (group G2) and pMPERTM/pHIV30 (group G3) at 12 days post-primary immunization. A stronger Tfh response was observed with pHIV30, as shown in Supplementary Fig. 5C, consistent with the findings presented in Figs. 5A and 5E (sG1 vs pG2). In a subsequent experiment, we also analyzed antigen-specific, isotype-switched memory B cells 30 days after priming with pMPERTM/sLACK (mG1) or pMPERTM/pHIV30 (mG2). The frequency and the number of MPER-specific memory B cells induced by pMPERTM/pHIV30 both trended larger than that by pMPERTM/sLACK (Fig. 6G). Likewise, the MPER-specific recall antibody response was much higher 10 days after the homologous boost with pMPERTM/pHIV30 compared to that with pMPERTM/sLACK (Fig. 6H). On the other hand, exposure of pHIV30 arrayed on the liposome surface induced higher anti-HIV30 antibody response compared to that induced by sHIV30 (Fig. 6I and Fig. 5D). Despite the greater the anti-HIV30 antibody titer than the anti-MPER antibody titer, the difference was smaller compared to that between anti-LACK and anti-MPER antibody titers (Fig. 6I).

## Discussion

In this study, we explored heterologous immunization strategies to understand factors that mitigate further expansion of existing off-target dominant B cell responses and potentiate subdominant MPER-specific B cell responses upon subsequent boosts. Earlier studies suggest that the relative affinity of the BCR determines whether a B cell will dominate the GC reaction^65,66^. In a noncompetitive environment, low affinity B cells can dominate the GC reaction ^67^, implying that immunogenicity of a MPER/liposome vaccine can be context dependent. Our results showed that the MPER-specific B cell responses are influenced by the competing B cell epitope directed against the helper peptides and their formulation since all other parameters such as antigen amount, valency and adjuvant are constant in the various liposome vaccines. The formulation methods of the helper peptides in the liposome vaccine appeared to modulate the amount of antigen for CD4 T cell help, influencing the GC responses and the resulting antigen-specific antibody responses with respect to their magnitude and affinity maturation. This was mainly associated with enhancing encapsulation efficiency of sHIV30 and sOVA through palmityl linkage to lipid to deliver more equivalent amounts of the helper peptide to draining lymph nodes.

Although very weak HIV30- and OVA-specific antibody responses are elicited by NpMPER/ sHIV30 and NpMPER/sOVA, the weak GC responses induced by the two helper peptides compared to the sLACK make it difficult to evaluate intrinsic B cell affinity for each helper peptide. Given the similar efficacy of CD4 T cell help on the GC responses induced by pLACK, pHIV30 and pOVA, respectively, the greater serum IgG response for the LACK peptide likely implies an intrinsically greater affinity/avidity of B cells for LACK. The data in Fig. 5E-F and 5H regarding LACK relative to the HIV30 and the OVA helper peptides supports this notion. Interestingly, MPER-specific GC B cells induced after priming and subsequent serum IgG responses were lowest with NpMPER/pLACK compared to pHIV30 and pOVA. Accordingly, difference in the magnitude of the vaccine-elicited antibodies specific for the MPER vs LACK was larger than that for the MPER vs pHIV30 and for the MPER vs pOVA, indicating the influence of the helper peptide B cell epitopes on the MPER immune responses in the context of the liposome vaccines.

In heterologous prime-boost immunization settings (Fig. 2A), dominant sLACK-specific B cells likely suppressed pMPERTM-specific B cells through interclonal competition for limited CD4 T cell help during GC reactions after priming. Subsequent boosts that replaced LACK with sHIV30 and sOVA peptides significantly blunted LACK-specific recall responses (Fig. 4E and Supplementary Fig. 2C). This reduction in competition allowed for greater clonal expansion and enhanced plasma responses from MPER-specific B cells. By dampening the activation of pre-existing memory B cells targeting LACK and HIV30 during sequential boosts with heterologous helper peptides, MPER-specific memory B cells—having lower activation thresholds than naïve B cells^68,69^—may be better supported.

Additionally, the relative affinity differences between NpMPER and HIV30 or OVA were smaller compared to NpMPER vs. LACK, possibly contributing to the improved response. The narrower affinity gap in the boosts using NpMPER/sHIV30 followed by NpW680A/sOVA may have facilitated quicker antigen access for MPER-specific memory B cells and improved recruitment of both memory and naïve B cells into secondary GCs, ultimately enhancing MPER-specific responses.

Overall, the combination of a reduced affinity gap and diminished activation of helper peptide-specific memory B cells appears to offset the lower CD4 T cell help provided by sHIV30 and sOVA compared to sLACK (Fig. 2A, G4 regimen). In contrast, the homologous helper peptide boosting strategy using NpMPER/sLACK followed by NpW680A/sLACK (G3 regimen) led to expansion of MPER-specific B cells but produced weaker antibody responses and no improvement in affinity. This occurred despite greater CD4 T cell help, indicating that in these heterologous immunization settings, affinity gap between MPER- and helper peptide-specific B cells plays a more critical role than the absolute amount of CD4 T cell help provided. High-affinity B cells thus appear to have a competitive advantage in accessing limited T cell help.

B cells presenting high surface peptide/MHC complex density are preferentially recruited into early GCs^70,71^. Similarly, increasing the quantity of HIV Env-specific CD4 T cell help improved the recruitment of rare VRC01 bnAb precursor B cells ^72^. Our results in Fig. 6 show that increasing CD4 T cell help following subsequent boosts with pHIV30 and pOVA supports greater affinity gain for MPER-specific serum IgG antibodies compared to the boosts with sHIV30 and sOVA (Fig. 6E). However, the priming immunogen exerted greater influence on the outcome of MPER-specific responses, affecting the magnitude and the affinity maturation of the MPER-specific serum IgG responses. This MPER augmentation was associated with increased T cell help provided by pHIV30 over sLACK and the reduced affinity gap between B cells specific for the pMPERTM and for the pHIV30. This contrasts with the large gap observed for MPER-versus LACK-specific B cells in GCs. In turn, the closer parity resulted in greater MPER-specific memory B cell generation and anti-MPER antibodies with higher affinity at 10 days after homologous booster immunization with pMPERTM and pHIV30. Furthermore, the magnitude of MPER-specific antibody responses was also positively correlated with the induction of functionally relevant antibodies capable of binding ADAgp145 (Fig. 6F). Given the similar helper function and antibody affinity gap between OVA vs MPER and HIV30 vs MPER (Fig. 5E-F and 5H), priming with pMPERTM/pOVA would have resulted in the similar outcome as the one induced by pMPERTM/pHIV30 priming. However, we cannot exclude the possibility that following boost with NpMPER/sLACK instead of NpMPER/pHIV30 after the priming with pMPERTM/pOVA may have limited MPER-specific recall responses in view of the immunodominant response directed at the LACK.

Consistent with our findings, priming with HIV-1 fusion peptides conjugated to keyhole limpet hemocyanin (KLH) proved more effective than an HIV-1 Env trimer vaccine in eliciting cross-clade neutralizing antibodies^73^, highlighting the critical role of the priming immunogen. Insufficient priming of MPER-specific B cells may limit the ability to generate a broad recall response following heterologous boosting.

Although the heterologous immunization regimen (Fig. 6A, G3) increased gp145 binding in polyclonal antibodies, no serum neutralizing activity was observed. The absence of neutralization may be due to insufficient serum antibody concentration capable of gp145 binding, lack of proper affinity maturation and/or differences in CDR3H length and lipid binding ability of the antibodies elicited from mice compared to human bnAbs. Alternatively, because MPER/liposomes do not mimic viral structural constraints, they may prompt antibodies with a suboptimal approach angle compared to that of human bnAbs. Future characterization of recombinant monoclonal antibodies is necessary to investigate these factors.

Our previous work demonstrated that repeated immunizations with pMPERTM/sLACK elicited high titer antibodies targeting the MPER. Paradoxically, these antibodies were unable to bind surface-expressed ADA gp145^49^. X-ray crystallography of vaccine-induced recombinant mAbs (rmAbs) revealed structural similarities to human bnAbs, but their approach angles were sterically incompatible with MPER access on the virion. These findings are corroborated by recent studies involving structural analysis of antibodies elicited by germline-targeting MPER-scaffolded immunogens ^74^ collectively underscoring the importance of MPER immunogen designs that incorporate structural features mimicking the steric hindrance of the viral membrane and the Env trimer ectodomain. Such designs may more effectively select for MPER-specific antibodies with favorable angles of approach for gp160 trimer engagement, increasing the potential for both higher serum titers and neutralizing activity. Nonetheless, as revealed herein, MPER immunogenicity is shaped by a multitude of variables, including not just immunogen structure but also the formulation, delivery, and immunological context of the vaccine. The development and affinity maturation of MPER-specific B cell responses remain particularly challenging due to their subdominant nature. Thus, while an optimized immunogen is necessary, it is unlikely to be sufficient alone. Our current study highlights that optimal immunogen design, while crucial, must be coupled with strategies that modulate immune hierarchies and promote MPER-focused responses. These insights provide an important foundation for refining both immunogen design and vaccination strategies.

In our heterologous MPER/liposome vaccination approach, B cell responses directed at dominant helper epitopes can be mitigated by replacing helper peptides with each booster immunization. This strategy is readily achievable with the peptide/liposome vaccine platform, offering a level of flexibility not possible with complex protein immunogens. Importantly, our findings suggest a potential strategy to suppress dominant off-target epitope-specific immune responses induced by prior vaccinations, thereby addressing the challenge of antigenic sin commonly seen in HIV vaccination. Introducing greater sequence variation in off-target epitopes with each booster, combined with enhanced affinity for target epitopes in heterologous immunogens, may improve immune focusing and overall vaccine efficacy.

## Methods

### Immunization

All animal experiments were carried out in accordance with procedures and protocols approved by the Dana-Farber Cancer Institute (DFCI) and Harvard Medical School Animal Care and Use Committee Institutional Review Board. Six-eight-week-old Balb/c mice were purchased from Taconic Bioscience (Germantown NY) and were housed and maintained in a specific pathogen free facility at DFCI. Mice were immunized subcutaneously with 50 µL of MPER/liposome vaccine per side, as described in the following section. Blood samples were collected via tail vein puncture at various time points throughout the study. At the study’s conclusion, mice were euthanized, and lymph nodes and spleens were harvested for further analysis.

### MPER/Liposome Preparation

Liposomes were prepared according to methods described previously ^59^. Briefly, N-terminally palmitoylated peptides including pMPERTM (palm-DLLELDKWASLWNWFNITNWLWYI KLFIMIVGGLV-CONH2), NpMPER (palm-DLLELDKWASLWNWFNITNWLWYIK-CONH2), or NpW680A (palm-DLLELDKWASLWNWFNITNWLAYIK-CONH2) were combined with lipids 1,2-dioleoyl-sn-glycero-3-phosphocholine (DOPC) (Avanti Polar Lipids Inc., 850375), 1,2-dimyristoyl-sn-glycero-3-phosphocholine (DMPC) (Avanti Polar Lipids Inc., 850345), 1,2-dioleoyl-sn-glycero-3-phospho-(1′-rac-glycerol) (DOPG) (Avanti Polar Lipids Inc., 840475), and 1,2-distearoyl-sn-glycero-3-phosphoethanolamine-N-[methoxy(polyethyleneglycol)-2000] (DSPE-PEG) (Avanti Polar Lipids Inc., 880120) at a molar ratio of 2:2:1:1. Additionally, N-terminally palmitoylated helper peptides pLACK, pHIV30, or pOVA were also included in the formulation of these liposomes (1:200 peptide to lipid ratio). The mixture was dried under a nitrogen stream and kept under vacuum overnight. The following day, liposomes were rehydrated in 1X Phosphate Buffered Saline (PBS), followed by six rounds of vortexing for 30 seconds at five-minute intervals. This was followed by six freeze-thaw cycles: the mixture was snap-frozen in liquid nitrogen for up to a minute and thawed at 37°C for five minutes. Finally, the liposomes were extruded 21 times through a 200 nm polycarbonate membrane (Whatman Inc., WHA10417004) to ensure uniform size. For MPER/liposome vaccines containing soluble helper peptides, a final concentration of 40 µg per mouse of soluble peptides sLACK, sHIV30, or sOVA were added to the MPER/liposomes at the time of rehydration in PBS. Cyclic dinucleotide (CDN) adjuvant-containing liposomes were prepared by separately drying the lipids as above and incorporating the CDN for encapsulation at the time of rehydration to facilitate its encapsulation, followed by the same vortexing and freeze-thaw process. After extrusion, CDN liposomes were subjected to ultracentrifugation at 25PSI for 2 hours. The resulting pellet was resuspended in 1X PBS, and CDN encapsulation efficiency was quantified using a spectrophotometer. The final CDN concentration in the vaccine formulation was 5 µg per mouse, which was mixed with MPER/liposome vaccines at the time of immunization. The pMPERTM/liposome was prepared at a molar peptide-to-lipid ratio of 1:300, while the NpMPER/liposome was at 1:200. N-terminally palmitoylated helper peptides were consistently used at a 1:200 ratio in all experiments. The total lipid concentration was 25 mg/mL, with each mouse receiving 2.5 mg in a 100 µL volume. Mice (N=5 per group, unless otherwise indicated in the figure legends) were immunized subcutaneously on day 0 (primary immunization), followed by a booster on day 30 and a second booster 30 days after the first, unless specified otherwise in the figure legends.

### Enzyme-linked immunosorbent assay (ELISA) to quantify mice serum IgG

Antigen-specific IgG responses in mice sera was quantified using ELISA as described ^49^. Ninety-six well Immulon 2HB plates (Thermo Scientific, 3455) were coated overnight at 4°C with 100 μl of DOPC/DOPG (4:1) arraying NpMPER, pLACK, pHIV30 or pOVA at 1:50 or 1:500 peptide-to-lipid ratio (final concentration of 100 μg/ml liposome in PBS). The following day, plates were washed three times with 0.1% BSA-PBS and blocked with 100 μl of 1% BSA-PBS per well for 4 hours at RT. Mouse sera or polyclonal IgG antibodies, starting a dilution of 1:500, in 1% BSA-PBS were incubated overnight with gentle rocking at 4°C. After washing, specific binding was detected using goat anti-mouse immunoglobulin (IgG) secondary antibody tagged with horse radish peroxidase (HRP) (Bio-Rad, 1706516, 1:2000) for 1 hour at 4°C. Further, the plates were washed twice with 0.1% BSA-PBS and twice with PBS. Bound antibody was detected by incubating with o-phenylenediamine (OPD) (Sigma-Aldrich, p9029) solution in citrate buffer (pH 4.5) for 7 minutes. The OPD reaction was stopped with 2.25 M H₂SO₄, and absorbance was measured at 490 nm using a Victor X4 plate reader (Perkin-Elmer).

### Calculation of % relative binding affinity by ELISA

First, to determine how MPER peptide density and antibody concentration affect antibody binding trends and % relative affinity, we compared ELISA titers of purified polyclonal antibody response against NpMPER/liposome at peptide to lipid ratio of 1:50, 1:250, 1:500, 1:1000, and 1:2000, respectively, as a reference. For each ratio lower than 1 to 50, we calculated the percent relative affinity by normalizing the optical density (OD) to the corresponding OD value from the peptide to lipid ratio of 1 to 50. When graphs of percent relative affinity are plotted against antibody dilution, linear curves are observed in a certain range for peptide-to-lipid ratios of 1:500 and 1:1000, leading to more reliable prediction. Ultimately, the 1:500 peptide-to-lipid ratio was selected for the relative affinity calculation because it provided a more dependable linear curve range, especially when immune response levels were low.

The % relative affinity in the following experiments was then calculated as the ratio of OD_490_ values obtained at low antigen density (1:500) to those at high antigen density (1 to 50) for the same serum dilution. To minimize signal saturation and reduce assay variability, only OD values below 3.0 and above 0.1 were included in the analysis. In practice, the OD values corresponding to approximately two dilutions below the saturation point were used to calculate the ratio, and the average of the two corresponding dilution points was taken as the final relative affinity value. This value was then multiplied by 100 to express the results as percent relative affinity for data visualization and comparison. For comparisons between experimental groups, the same dilution point selected for one group was consistently applied to the corresponding comparison group to ensure methodological consistency. Data was analyzed using GraphPad prism v10.

### ELISPOT quantification of antigen specific ASCs

Antigen-specific antibody secreting cells (ASCs) from bone marrow and spleen of Balb/c mice were quantified by Enzyme-linked immunosorbent spot (ELISPOT) as described earlier ^75^. Assay was performed in 96-well 0.45-μm, hydrophobic, high-protein-binding Immobilon-P polyvinylidene difluoride (PVDF) membrane plates (EMD Millipore, Billerica, MA). Plates were activated with 35% ethanol for 30 sec and then washed eight times with distilled water. The activated plates were incubated with 100 μl per well of 100μg/ml NpMPER or MPER-N or MPER-C liposome formulations in PBS (1:50 or 1:500 peptide: lipid ratio; 4:1 DOPC:DOPG ratio) overnight at 4°C. Next day, the plates were washed six times with 0.1% BSA-PBS, blocked with 200 μl/well 1% BSA-PBS for 4h at RT. Plates were briefly washed with RPMI 1640 medium supplemented with 10% FBS, glutamine, 2-mercaptoethanol, and penicillin-streptomycin, and then blocked for 1 h at 37°C with the same medium. Bone marrow and spleens were isolated from mice and single cell suspensions were obtained by filtration through 70 μm cell strainers in growth media. The cell suspensions were strained again through 70 μm, and 500,000 cells were added to the wells in 50-μl volumes. Cells were incubated overnight at 37°C and 5% CO₂ in a humidified chamber. The next day, wells were washed six times with 0.1% BSA-PBS and blocked for 1.5 hours with 1% BSA-PBS. After discarding the blocking buffer, bound antibody was detected using 0.6 μg/ml alkaline phosphatase (AP)- conjugated goat anti-mouse secondary anti-IgG antibody (Jackson ImmunoResearch, West Grove, PA) in 1% BSA-PBS. To visualize MPER-specific bound antibody, wells were washed eight times, and 100 μl of 5-bromo-4-chloro-3-indolyl phosphate/nitro blue tetrazolium (BCIP/NBT) solution was added for 5 minutes. Plates were washed with distilled water and dried overnight. Spots were quantified using a CTL ImmunoSpot ELISPOT plate reader and ImmunoSpot 3 software (CTL, Shaker Heights, OH).

### Surface plasmon resonance (SPR) measurements

BIAcore experiments were performed as described ^59^ using a BIAcore 3000 or BIAcore 1K system with either a Pioneer L1 sensor chip or a Series S L1 chip at 25°C. Data analysis was conducted using BIAevaluation 3.1 software (BIAcore) for BIAcore 3000. The peptide/liposome formulations (150–250 μM) in 20 mM HEPES, 0.15 M NaCl, pH 7.4 (HBS-N) were applied to the sensor chip surface at a flow rate of 5 μL/min to form a supported lipid bilayer. To remove multilamellar structures, 20 μL of 25 mM sodium hydroxide was injected at 100 μL/min, stabilizing the baseline response units (RU) between 3000 and 6000, corresponding to the immobilized bilayer. Polyclonal IgG antibodies were purified from mouse sera using a GammaBind Plus Sepharose column (GE Healthcare). Purified antibodies 30-40μg/ml were injected over the peptide-liposome complex at a flow rate of 20-50 μL/min for 3 minutes. Following binding analysis, the peptide-liposome complexes were removed using sequential injections of 40 mM CHAPS and a 6:4 mixture of 50 mM sodium hydroxide and isopropyl alcohol. The same protocol was used to test antibody reactivity against both MPER-N and MPER-C peptide/liposome complex as well as pMPERTM/liposome and NpW680A/liposome.

For epitope mapping, DOPC/DOPG liposomes (150–250 μM) were injected at 5 μL/min to establish a lipid bilayer on the sensor chip. To remove excess multilamellar structures, 20 μL of 25 mM sodium hydroxide was injected at 100 μL/min, resulting in a stable baseline (RU: 4500–6000). Alanine-scanning mutagenesis was performed on parental MPER sequences (HxB2) used for immunization. Peptide variants (0.5 μM) were freshly prepared in the running buffer before injection. Each peptide (60 μL) was injected over the immobilized liposome bilayer at 5 μL/min, followed by antibody injection (40–80 μg/mL) for 3 minutes at 10 μL/min. To regenerate the sensor surface, liposomes were removed using 25 μL of 40 mM CHAPS at 5 μL/min, followed by 10 μL of a NaOH (50 mM)/isopropyl alcohol (6:4) mixture at 20 μL/min. Each peptide injection was performed on a freshly prepared liposome surface. The relative reactivity of purified polyclonal IgG antibodies to MPER alanine mutants was determined by measuring response units at the 3-minute dissociation time point. Sensorgrams were recorded during a 3-minute association phase followed by a 3-minute dissociation phase. At least two independent experiments were conducted, varying peptide and antibody concentrations to assess the relative binding reactivity to each MPER mutant.

### ADA gp145 binding assay using Flow cytometry

HEK293T cells (ATCC, CRL-3216) were cultured in DMEM supplemented with 10% fetal bovine serum (FBS), 2 mM L-glutamine, and 1% penicillin-streptomycin. Cells, were transiently transfected with HIV-1 ADA gp145 Env at 60–70% confluence using polyethyleneimine (PEI Max, Polysciences) in a 1:3 plasmid-to-PEI (weight to weight) ratio as described ^49^. Cells were harvested 48 hours post-transfection in FACS buffer (1x PBS with 2% FBS, 1 mM EDTA, and 0.1% sodium azide). 100,000 cells were incubated with or without 5 μg polyclonal IgG antibodies at room temperature for 1 hour. Cells were washed 2X with FACS buffer followed by staining with Phycoerythrin (PE)-conjugated goat anti-mouse IgG (Southern Biotech, 1030-09, 1:500) and Zombie Aqua Fixable viability dye (BioLegend, 423101, 1:500) at 4°C for 30 minutes. Cell-surface fluorescence was analyzed using a BD LSR Fortessa. Data was analyzed using Flow Jo and median fluorescence intensity (MFI) was calculated by subtracting the signal from untransfected cells stained with the corresponding polyclonal IgG antibodies.

### Flow cytometry analysis

Single-cell suspensions were prepared from spleen tissue as described previously ^50^ by mashing the tissue through a 70 μm nylon mesh cell strainer (Corning, Durham, NC). Cells were then resuspended in FACS buffer. For each sample, 5 million cells were transferred to the wells of a conical-bottom 96-well plate and sequentially stained for 20 minutes on ice with Fixable Viability Zombie Aqua or Zombie NIR dyes (Biolegend), followed by washing with FACS buffer. Antibody staining was performed by first resuspending the cells in 50 μl of Fc block (CD16/32, BD Biosciences) and incubating on ice for 10 minutes. Afterward, cells were mixed with 50 μl of an antibody cocktail targeting T follicular helper (TfH) marker (CD4-Pacific blue (clone GK1.5), CD44-PECy7(clone IM7), CD62L-BV611(clone MEL-14), CXCR5-APC (clone L138D7), and PD1-FITC (clone 29F.1A12) (Biolegend). For antigen-specific GC B cell staining (MPER- and LACK, HIV30 or OVA-specific), cells were stained with viability dyes followed by incubation with 100 μg/ml NpMPER formulated with fluorescent 18:1 Liss Rhod PE and 1,2-dioleoyl-sn-glycero-3-phosphoethanolamine-N-(lissamine rhodamine B sulfonyl) (ammonium salt) along with pLACK, pHIV30 or pOVA formulated with fluorescent 18:1 PE CF1,2-dioleoyl-sn-glycero-3-phosphoethanolamine-N-(carboxyfluorescein) (ammonium salt) liposomes, respectively (Avanti Polar Lipids, Inc, AL, USA). This incubation was performed for 1 hour on ice. After staining, the cells were washed twice with 200 μl of FACS buffer, followed by a Fc block as described above. Cells were then stained with antibodies targeting GC B cell markers (B220-PerCP/BV85 (clone RA3-6B2), IgD-BV510/APCCy7 (clone 1126C2A), CD38-BV421 (clone 90), GL7-Pac-Blue (clone GL7), IgG-APC (clone Poly4053) (Biolegend) and analyzed on an LSRFortessa II cell analyzer (BD Biosciences). Data analysis was performed using FACS Diva and FlowJo software (version 10.8.1, Tree Star). Cell gating strategies are outlined in Supplementary Figs. 1B and 3. Flow cytometry was conducted at the Dana-Farber Cancer Institute.

### Ex Vivo restimulation of antigen-specific CD4⁺ T Cells

To evaluate antigen-specific CD4⁺ T cell responses following immunization, splenocytes were isolated from mice immunized with MPER/liposomes containing pLACK, pHIV30, or pOVA peptides, boosted on day 30, and harvested on day 40. Antigen-specific responses were assessed using intracellular cytokine staining (ICS), activation-induced marker (AIM) assay, and IFN-γ ELISpot.

#### Intracellular Cytokine Staining (ICS) Assay

Spleens were harvested from immunized mice and splenocytes were isolated by mechanical dissociation followed by red blood cell lysis using ACK lysis buffer. For in vitro stimulation, 1×10^6^ splenocytes were plated in 96-well round-bottom plates and incubated with 5 µg/mL of peptide (LACK, HIV30, or OVA) for 2 hours at 37°C in complete RPMI (10% FBS, 1% Pen/Strep, 1% L-glutamine, 1 mM sodium pyruvate, 0.1 mM non-essential amino acids, 50 µM β-mercaptoethanol). After 2 hours, Brefeldin A (BD GolgiPlug) was added to a final concentration of 10 µl/mL, and cells were incubated for an additional 4 hours (total 6 hours of stimulation).

Following stimulation, cells were washed with PBS and stained with a fixable viability dye (Zombie NIR Biolegend) and surface markers anti-CD3 (clone 17A2), CD4 (clone GK1.5), CD44 (clone IM7) for 30 minutes at 4°C. After washing, cells were fixed and permeabilized using BD Cytofix/Cytoperm buffer according to the manufacturer’s instructions. Intracellular staining was performed with antibodies against IFN-γ (clone XMG1.2) in perm/wash buffer for 30 minutes at 4°C. Samples were acquired on a BD LSRFortessa flow cytometer and analyzed using FlowJo software (BD Biosciences). Gating strategy involved initial exclusion of dead cells, followed by selection of CD3⁺CD4⁺ T cells, and analysis of CD44^+^IFN-γ expression to determine antigen-specific T cell responses.

#### Activation-Induced Marker (AIM) Assay

Antigen-specific CD4⁺ Tfh cell responses were assessed using an AIM assay. 1×10^6^ Splenocytes were isolated from immunized mice and plated in 96-well round-bottom plates and stimulated in vitro with 5 µg/mL of peptide (LACK, HIV30, or OVA) in complete RPMI media for 18 hours at 37°C. No Brefeldin A was used during the stimulation. Following incubation, cells were stained with fixable viability dye (Zombie NIR) and fluorochrome-conjugated antibodies against CD3, CD4, CD44, CXCR5 (L138D7), PD-1 (29F.1A12), CD25 (PC61), and OX40 (OX-86). Staining was performed in FACS buffer (PBS + 2% FBS) for 30 minutes at 4°C. Stained cells were acquired on a BD LSR Fortessa flow cytometer and analyzed using FlowJo software (BD Biosciences). The gating strategy included exclusion of doublets and dead cells, followed by identification of CD3⁺CD4⁺CD44⁺ T cells. Within this population, Tfh cells were defined as CXCR5⁺PD-1⁺. Antigen-specific activation was quantified by identifying co-expression of CD25 and OX40 within the Tfh (CXCR5⁺PD-1⁺) gate. Fluorescence minus one (FMO) control were used for gating to ensure accurate identification of AIM⁺ cells.

#### IFN-γ ELISpot Assay

To assess antigen-specific CD4⁺ T cell responses, splenocytes (1 × 10⁵ cells/well) were plated in triplicates on anti–IFN-γ–coated ELISPOT plates and stimulated with 5 µg/mL of LACK, HIV30, or OVA peptides for 18 hours. Unstimulated wells served as negative controls. After incubation, plates were washed and incubated with biotinylated anti–IFN-γ detection antibody. Subsequent detection steps were performed as described previously.

### Statistical analysis

All mouse experiments were conducted two to three times, with each group consisting of 4–6 mice, unless indicated in the figure legends. Data were analyzed using either a non-parametric t-test for comparisons between two datasets or Kruskal-Wallis test with Dunn’s comparison test for multiple groups. For multiple group analyses, two-way ANOVA (or a mixed-effects model) was used for row vs. column comparisons, with Sidak’s or Tukey’s multiple comparison tests applied as appropriate. Statistical differences were denoted by p-values: ****p < 0.0001, ***p < 0.001, **p < 0.01, *p < 0.05, while p > 0.05 was considered not significant. Analyses were conducted using GraphPad Prism v10.

### Lead Contact

Further information and requests for resources and reagents should be directed to and will be fulfilled by the lead contact, Mikyung Kim.

### Materials availability

All cell lines, plasmids, and other stable reagents generated in this paper are available from the lead contact under a complete Materials Transfer Agreement.

### Data and code availability

- This paper does not report original code.
- Any additional information required to reanalyze the data reported in this paper is available from the lead contact upon request

## Acknowledgments

This work was supported by National Institutes of Health Grant AI145509 to M.K.; AI181597 to E.L.R. and M.K.; AI126901 to E.L.R.

## Supplementary Information

**Supplementary Figs. 1-5**

