## Supplementary Fig for "Heterologous immunization modulates B-cell epitope competition between helper peptides and the MPER segment in MPER/liposome vaccines"

**Supplementary Information**

**Supplementary Fig. 1 Epitope mapping of anti-MPER antibodies elicited by NpMPER/liposome and MPERTM/liposome vaccines and comparative analysis of primary GC B cell responses induced by pMPERTM/liposome and NpMPER/liposome vaccines**

A) Epitope Specificity of MPER-specific polyclonal IgG induced by NpMPER/Liposome vs. MPERTM/liposome immunization. BALB/c mice (n = 5 per group) were immunized with either NpMPER/liposome or MPERTM/liposome formulations, each containing sLACK helper peptides and a CDN adjuvant. Immune sera were collected 30 days following the third homologous immunization and subsequently purified for analysis. Data presented are adapted from a previously published study by our group. B) Representative flowcytometry gating strategy for identification of antigen specific GC B cells. C) GC B cell frequency, and total cell number from mice spleen collected at day 8 of immunization. Data is representative of one of the two independent experiments (n=6). Error bars represent mean ± SEM.

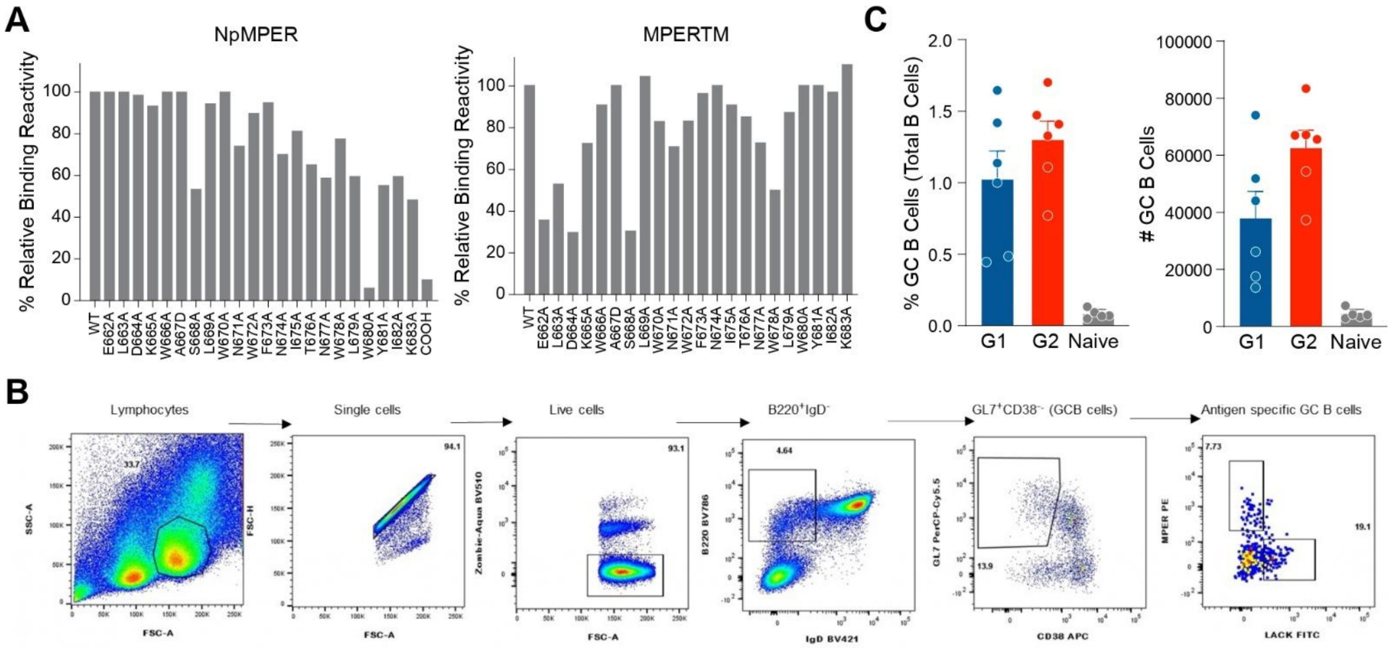

**Supplementary Fig. 2 Immunogenicity of various MPER/liposome vaccines**

A) Distinct epitope specificity of MPER-specific antibodies elicited by NpMPER, pMPERTM, and NpW680A immunogens, as measured by SPR analysis. Binding to alanine-substituted mutants was normalized relative to the wild-type sequence to reveal differences in epitope recognition. Three different groups of Balb/c mice were immunized s.c. three times with NpMPER/liposome, pMPERTM/liposome and NpW680A/liposome vaccines containing sLACK helper peptides and CDN adjuvant, respectively. Immune sera were collected 30 days after the final immunization and purified polyclonal IgG antibodies were used for the epitope mapping analysis. B) ELISA binding curves corresponding to Fig. 2B. C) ELISA titers against LACK, HIV30, and OVA antigens corresponding to each immunization regimen presented in Fig. 2A. Antigen-specific IgG responses were evaluated using polyclonal antibodies purified from immune sera. Data are presented as binding curves, with log-transformed serial antibody dilutions on the x-axis and absorbance values on the y-axis. Results represent one of two independent experiments. Error bars indicate mean ± SEM.

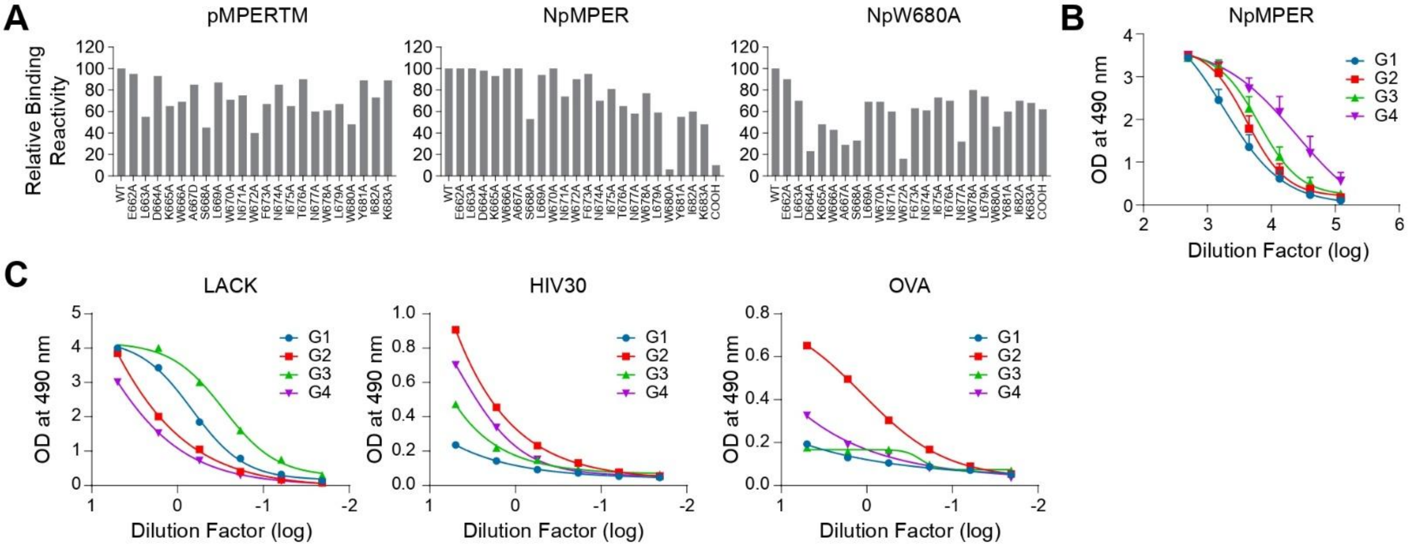

**Supplementary Fig. 3 Flow cytometry gating strategy for the identification of Tfh cells**

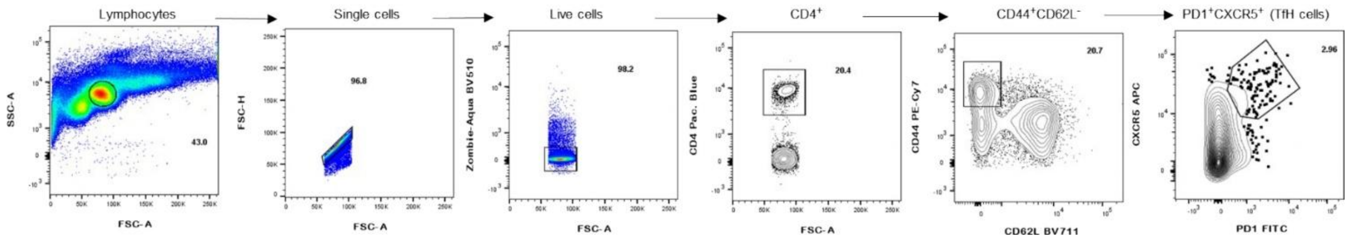

**Supplementary Fig. 4 Ex Vivo restimulation of antigen-specific CD4$^+$ T Cells**

Assessment of antigen-specific CD4$^+$ T cell responses following immunization was performed using splenocytes isolated from mice immunized with MPER/liposomes containing pLACK, pHIV30, or pOVA peptides, boosted on day 30, and harvested on day 40. A) Frequency of antigen-specific IFN-γ–secreting cells in total splenocytes determined by ELISPOT assay. (B) IFN-γ production by activated (CD44$^+$) CD4$^+$ T cells measured by intracellular cytokine staining. (C) Frequency of total Tfh cells (CD44+CXCR5$^+$PD-1$^+$) following peptide restimulation. (D) Antigen-specific Tfh activation measured by upregulation of CD25 and OX40 within the Tfh compartment. Statistically significant differences between different groups were determined Kruskal-Wallis test with Dunn's comparison test denoted by p values: *$p < 0.05$. Data are representative of two independent experiments with 5 mice per group. Error bars represent mean ± SEM.

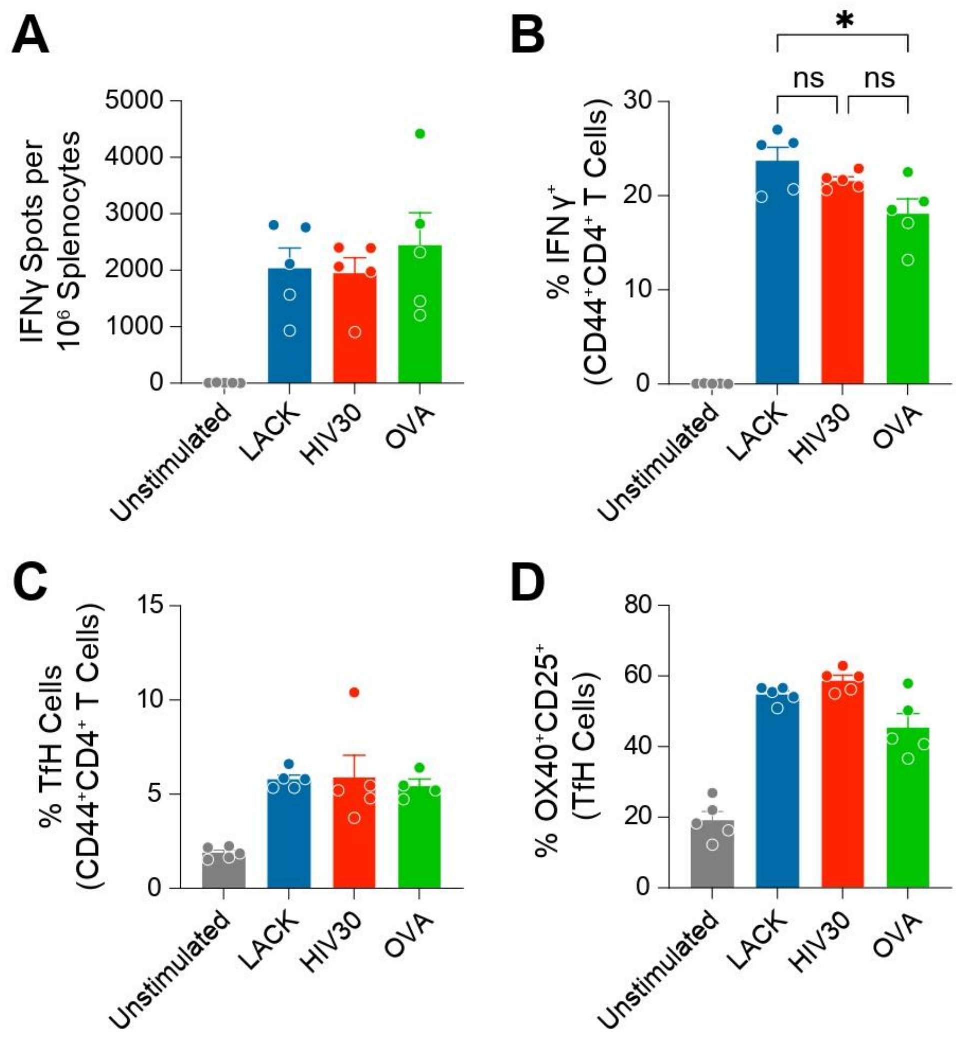

**Supplementary Fig. 5 Frequency and total number of Tfh cells induced by sLACK vs pHIV30**

Tfh cell frequencies were assessed in mice immunized with pMPERTM/liposome with sLACK or pHIV30 at 12 days post-primary immunization. Bars represent % Tfh cells from total CD4 T cells. Statistically significant differences between the two groups were determined using the Mann–Whitney test denoted by p values: *$p < 0.05$. Data are representative of two independent experiments with 5 mice per group. Error bars represent mean ± S EM.

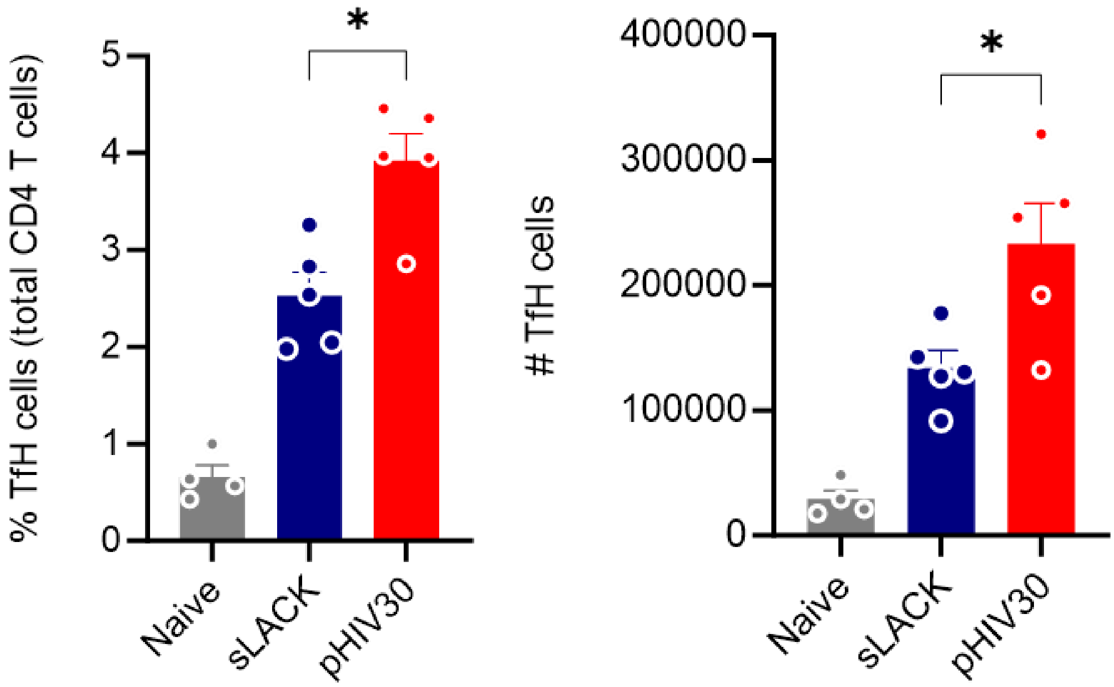
